# Flagellotropic bacteriophage selects for evolutionary remodeling of cell motility and chemotaxis in *Escherichia coli*

**DOI:** 10.1101/2025.05.06.652435

**Authors:** Jyot D. Antani, Austen Theroux, Fotios Avgidis, Jackson A. Wood, Birgit E. Scharf, Thierry Emonet, Paul E. Turner

## Abstract

Bacteria can evolve resistance to bacteriophage attack via mutations that alter phage receptors on the cell surface. Flagellotropic phages bind to motility-enabling flagella, creating a potential evolutionary trade-off, since losing flagellar function can reduce fitness in many environments. Phage-resistance studies typically characterize this trade-off using population-mean phenotypes, but selection can act on, and reshape, the underlying single-cell distribution of a trait in ways that mean values alone do not capture. Here, we exploit the well-developed and experimentally tractable model system of *Escherichia coli*–phage χ (chi) to ask how evolved resistance to a flagellotropic phage reshapes single-cell motility phenotypes, not just their population averages. We experimentally evolved *E. coli* populations under selection by χ, on soft-agar swim plates which favor motility. Whole genome sequencing revealed early emergence of non-motile mutants with disrupted motility genes, followed by motile, χ-resistant mutants carrying flagellin mutations. Swim-plate assays and single-cell tracking revealed heterogeneous motility outcomes among resistant lineages: some recovered or exceeded ancestor-level motility while others swam slower than ancestors, demonstrating that trade-offs and trade-ups can both arise from selection by the same phage—an outcome rarely captured in a single phage-resistance study. Single-cell tracking further showed that resistant lineages evolved increased tumble bias and altered swimming speed distributions, revealing shifts in trait heterogeneity invisible to mean-based measures. At the ensemble level, resistant mutants also showed remodeled kinase-response output in the upstream chemotaxis signaling pathway, indicating that resistance reshapes motility behavior both through receptor-level and signaling-level adaptation. Together, these findings demonstrate that evolved phage-resistance reshapes the receptor-linked (motility) phenotype across multiple phenotypic scales: from single-cell behavioral distributions to signaling network output, producing diverse and previously undocumented evolutionary outcomes under combined selection pressures.

**Author Summary:** Bacteria often rely on flagella, tail-like appendages that enable swimming, to move toward nutrients and escape undesirable environments. Many viruses that infect and kill bacteria (lytic bacteriophages) exploit these motility-enabling flagella to attach to bacterial cells, creating an evolutionary dilemma: a bacterium can become resistant by losing its flagella, but this sacrifices its ability to move. We used experimental evolution to study how motile bacteria respond to attack by such phages. By exposing *E. coli* to the flagellum-targeting phage χ over many generations, we found that resistance evolved in two stages. Bacteria first became resistant by losing motility entirely, but soon after, motile χ-resistant variants emerged carrying mutations in flagellin, the flagellum’s main building block. Some of these motile-resistant lineages swam even better than their ancestor, while others swam worse, revealing diverse evolutionary solutions to the same selective pressure. Using fluorescence-based signaling measurements, we found that resistant bacteria also altered the internal chemical signaling that controls swimming behavior, not just the physical structure targeted by the phage. These findings reveal how pathogens can retain or even enhance mobility while evading phage attack, with implications for designing phage-based therapies to treat infections by motile pathogens.

## Introduction

Bacteriophages (phages), the viruses of bacteria, have existed alongside their bacterial hosts for perhaps billions of years. Following infection and intracellular replication of bacteria, lytic phages destroy (lyse) host cells to release virus progeny, thereby imposing strong selection for bacterial populations to evolve defenses against phage attack [1,2]. Bacteria can evolve phage-resistance by mutating or suppressing the phage binding receptor(s) on the cell surface, because viruses use the structure(s) to initiate infection. Phages have evolved to exploit various exposed structures on host cell surfaces for attachment, including transmembrane channels, saccharides, and appendages. Binding to appendages that extend well beyond the cell surface may increase the probability of freely-diffusing phages to collide with such structures on host cells suitable for infection [3,4].

Flagellotropic phages infect bacteria by first interacting with the bacterial flagellum [5,6]. Flagella are threadlike extracellular appendages that sometimes extend up to multiple cell lengths. Rotation of flagella enables planktonic motility of bacteria in aqueous media as well as their swarming motility on semisolid surfaces [7,8]. Obstruction of flagellar rotation is thought to be the earliest signal in surface sensing and biofilm establishment by bacterial cells [9–11]. Flagellar motility allows bacteria to efficiently explore their environments through chemotaxis (directed movement along chemical gradients) [12], and differing motility can determine relative fitness (reproduction success) of bacterial variants (genotypes) in a population when movement is essential for obtaining limiting nutrients [13,14] and is considered a virulence factor for certain bacterial pathogens [15]. If mutations arise in the motility apparatus of bacteria to evade attack by flagellotropic lytic phages, movement through the environment may be negatively affected, thus representing a crucial fitness trade-off suffered by bacteria that evolve phage resistance. While bacteria and flagellotropic phages are thought to coexist in nature [5,16], the mechanisms by which bacteria evolve resistance against such phages remain poorly understood.

Bacterial motility and chemotaxis in *Escherichia coli* have been extensively studied over the last five decades [8,17–19]. Cells express multiple flagella distributed across the cell, each with a transmembrane flagellar motor at its base. The left-handed flagella bundle together when rotating in their default counterclockwise (CCW) direction and propel the cell forward, resulting in straight ‘runs’. When one or more motors switch to clockwise (CW) rotation, the bundle breaks, and the cell reorients due to forces in different directions, resulting in a ‘tumble’. Cells perform a run-tumble random walk (**Fig 1**A), in which binding of the intracellular response regulator CheY-P to the flagellar motor biases the motor toward CW rotation (tumbles). This regulator is part of the chemotaxis two-component system, in which the kinase CheA, when activated, phosphorylates CheY. CheA is part of the receptor array, which can sense extracellular ligand concentrations and accordingly modulate CheA activity. Another chemotaxis enzyme, CheZ, acts as a phosphatase for CheY-P (**Fig 1**B).

**Fig 1.**
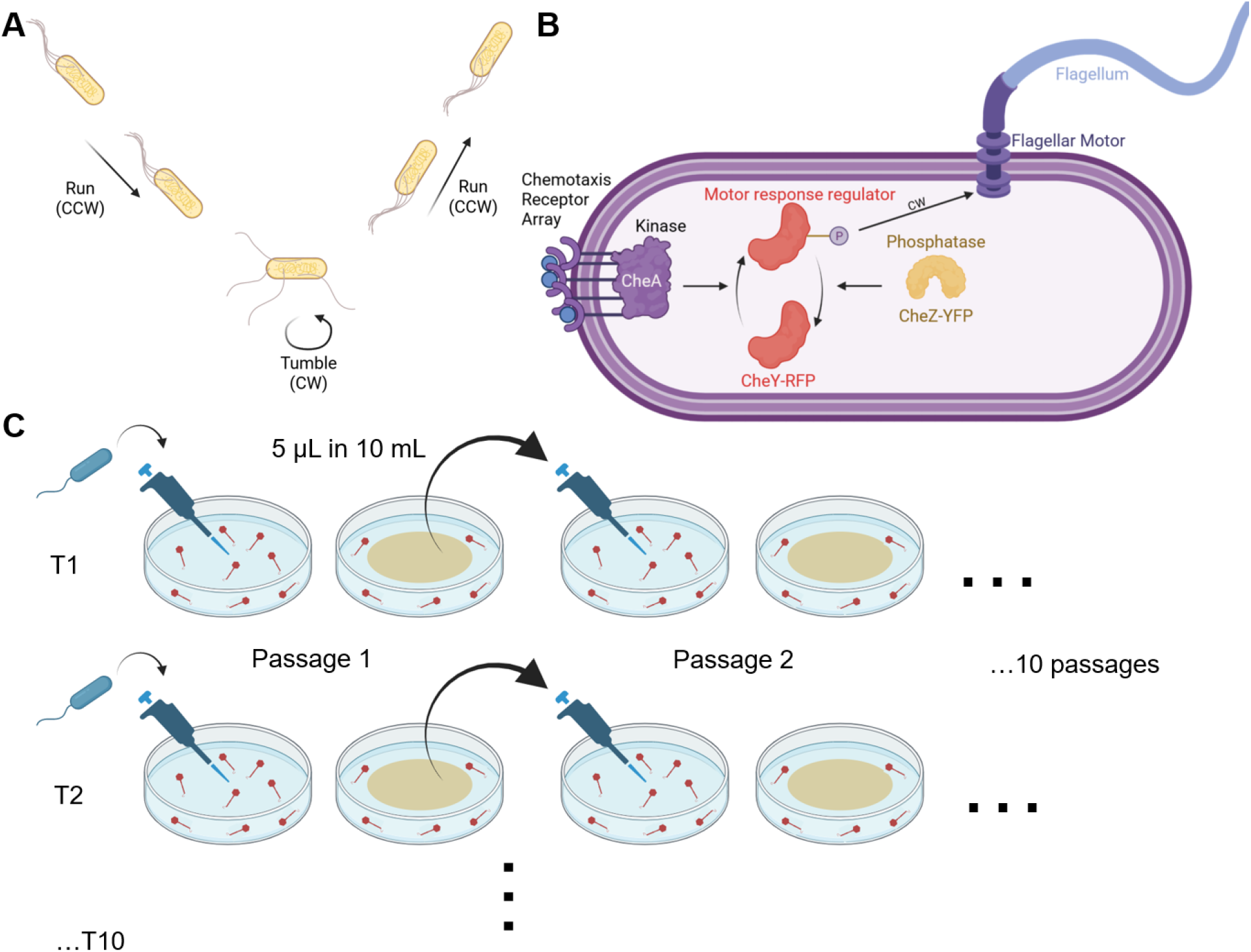
Background and Experimental Overview. **(A)** Flagella bundle together when rotating counterclockwise (CCW) and push the cell in a straight ‘run’. When one or more flagella turn clockwise (CW), the bundle breaks and the cell reorients, which is termed a ‘tumble’. Switching between runs and tumbles allows cells to navigate their environment. **(B)** The chemotaxis network controls the rotational direction of the flagellar motors through a two-component system. Receptors sense ligand molecules and accordingly adjust the kinase CheA’s activity. The activated kinase phosphorylates the response regulator CheY, while the phosphatase CheZ can dephosphorylate it. Förster Resonance Energy Transfer (FRET) between CheY-RFP and CheZ-YFP provides a readout of kinase activity. **(C) Swim-plate assay for evolution of bacteria against phage** χ. We inoculated bacteria into the center of swim plates (0.3% agar) embedded with phage χ. After 24-48 hours of incubation at 30°C (one passage), we transferred an aliquot to fresh plates containing phage χ. We continued this protocol for 10 passages, for 10 independently evolving replicate Treatment lineages T1-T10, and for four Control lineages C1-C4 evolved in absence of phage χ. Parts of this figure were created in BioRender. Turner, P. (2026) https://BioRender.com/51hbpmz.

Here, we used experimental evolution to examine whether populations of *E. coli* could maintain motility while evolving resistance against the virulent phage χ, a model flagellotropic phage. Phage χ requires fully functional, rotating flagella for successful infection [20,21]. While bacterial resistance to phages is known to involve receptor modification, little is known about how such modification reshapes the receptor-linked phenotype at the single-cell level across evolving populations. We chose the *E. coli*–x system specifically because *E. coli* motility and chemotaxis are exceptionally well-characterized and experimentally tractable at the single-cell level, allowing us to resolve not just whether resistance evolves, but how the underlying behavioral trait distribution is remodeled across a population during that process.

We conducted experimental evolution of bacterial populations founded by the motile *E. coli* strain MG1655 in replicated environments containing both phage selection pressure and positive selection for cell motility. Whole genome sequencing revealed that early in the experimental evolution, nonmotile χ-resistant mutants featured deletions in motility and chemotaxis genes. Whereas, later in evolution, phage-resistant mutants emerged which maintained motility and carried mutations in *fliC*, the gene encoding flagellin, which is the primary component of the flagellar filament. Across treatment populations, we observed parallel evolution of specific key mutations that impart single amino-acid substitutions in the outer D3 domain of flagellin. Finally, we assessed the evolutionary impact of key mutations on bacterial motility and chemotaxis using swim-plate assays, single-cell motility tracking, and FRET (Förster Resonance Energy Transfer)-based measurements of chemotaxis kinase activity [22,23] (**Fig 1**B), by comparing the ancestral strain to its evolved descendants, including control strains evolved in absence of phage χ. This study highlights the molecular adaptations and evolutionary outcomes experienced by populations of motile bacteria when selected to maintain motility while avoiding infection by a flagellotropic phage.

### Experimental Evolution Overview

Typical phage-bacteria coevolution experiments involve co-culturing phages and bacteria in shaking flasks with serial transfers (dilution bottlenecks into a flask containing fresh medium) performed every ∼24 h. Flagellar synthesis and operation are often energetically expensive: flagellar synthesis may account for ∼2% of the total cellular energy expenditure in model strains of *E. coli* and *Salmonella* and up to ∼10% in non-model bacteria [24,25]. When an anti-flagellar evolutionary pressure such as phage χ is present, bacterial mutants with defective flagellar synthesis may emerge as a simple adaptive strategy. In more natural and complex environments, however, bacteria must also maintain motility to locate nutrient-rich habitats. To simulate these conditions, we challenged *E. coli* to evolve under laboratory environments alongside phage χ while also selecting for bacterial motility. To do so, we introduced a high concentration of χ into ‘soft’ agar (0.3%), containing a tryptone-based growth medium that allows bacteria to move via chemotaxis-driven flagellar motility. We then inoculated bacteria from a freshly-grown culture of motile *E. coli* MG1655 (defined as ancestor or wild-type) in the center of the soft-agar plate and incubated it for up to 48 h at 30°C. The expectation was that motile bacteria would spread concentrically through the medium, expanding in population size to create a visible ‘swim’ ring, whereas non-motile bacteria would remain near the center of the agar plate. Serial passage (1:2000 population bottlenecking) was achieved by harvesting the plate contents, mixing 1:1 with liquid medium, and using a dilution of this mixture to repeat the process of inoculating the bacteria into the center of a fresh soft-agar plate containing χ. Samples from harvested plates were used to create frozen stocks that were stored for later analysis. This process was repeated for 10 passages total, for each of 10 independently evolving replicate lineages (Treatment lineages T1 through T10) descended from the original ancestor genotype (**Fig 1**C). This method favored evolution of χ-resistant bacteria that were additionally selected to maintain flagellar motility, which enables bacteria to access more nutrients. In addition, four control lineages (C1 through C4) were serially passaged identically, using plates where phage χ was absent.

## Results

### IS1-mediated deletions provide resistance to phage **X** in early evolved, nonmotile mutants

Early in the evolution experiment, we observed that bacteria in all 10 Treatment lineages (up to passage 2 in some lineages, up to passage 3 or 4 in others) failed to form a swim ring even after 48 hours of incubation. Instead, non-motile cells grew on the plate as isolated, ‘spotty’ colonies (**Fig 2**A). These bacterial populations appeared to evolve the expected (strongly selected) phenotype of inhibition of flagellar synthesis and/or rotation when passaged in an environment alongside phage χ; the flagellotropic phage is unable to bind and infect host cells that lack functional flagella [20,21] (**Fig 2**B). This phenotypic change in cell motility was not observed in the four Control lineages.

**Fig 2.**
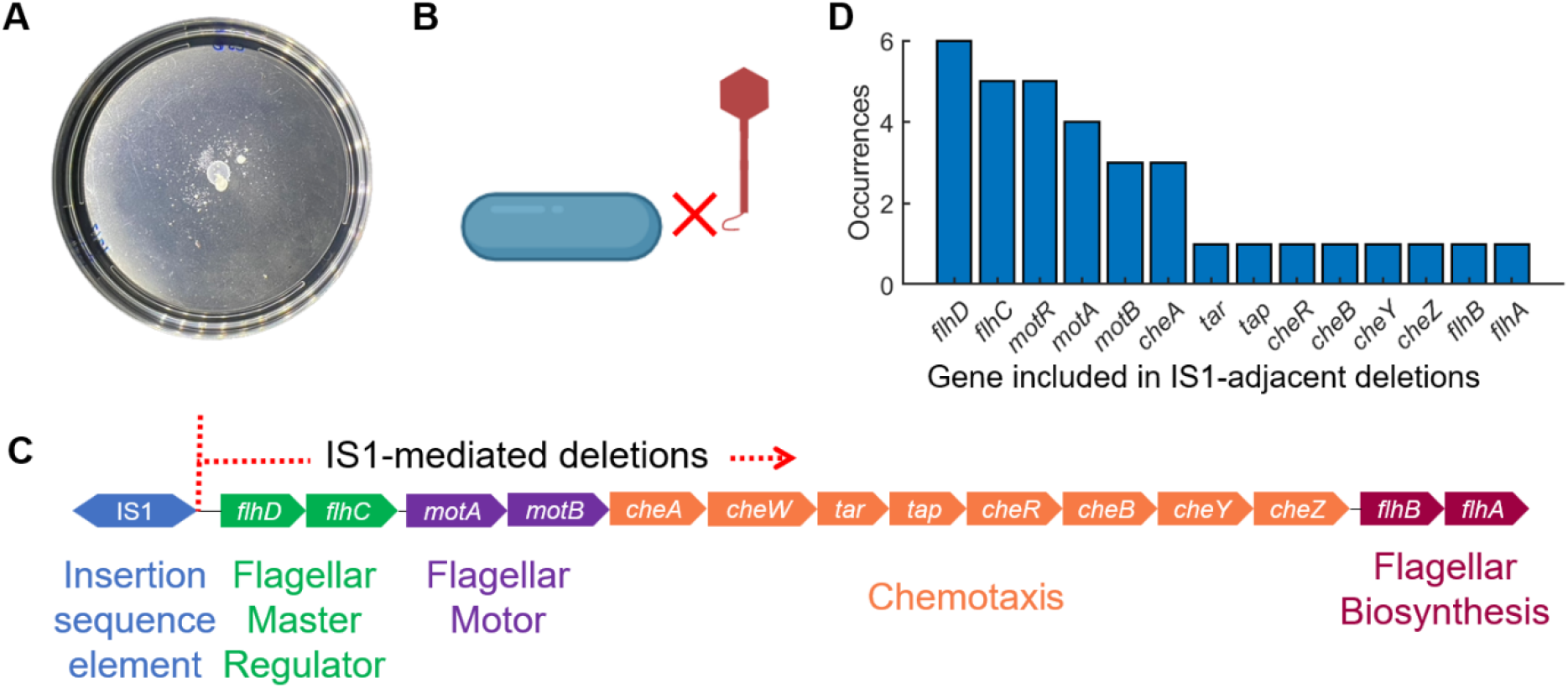
Early in evolution, selection tended to favorx-resistant mutants that were non-motile. **(A)** Early in experimental evolution (up to passages 2 and 3), the populations did not present visible swim rings and instead showed ‘spotty’ colonies of bacteria, likely started by low-frequency motile mutants. **(B)** Thus, these Treatment lineages developed resistance to phage χ by abolishing flagellar synthesis, as the virus cannot infect bacteria lacking functional flagella. **(C)** The motility-related mutations were mediated by gene deletions downstream of insertion sequence element IS1 (likely mediated by insertion of another IS1 copy), which is located in the upstream regulatory region of *flhD*. Genes removed by IS1-adjacent deletions included *flhDC*, which encode the flagellar master operon. The genomes of six representative nonmotile mutant populations were sequenced, and the overall frequency of deleted genes was determined. **(D)**. Parts of this figure were created in BioRender. Turner, P. (2026) https://BioRender.com/51hbpmz.

To probe the genetic basis of motility loss, we sequenced the population genomes of each Treatment lineage at passage 3 and analyzed the data with breseq [26]. Six of these lineages consisted of nonmotile χ-resistant bacteria (‘spotty’ colony morphologies), and we focused on these representative consensus genomes. We discovered that an insertion sequence element, IS1, mediated the disruption of motility-related genes. In the ancestral strain MG1655, this transposon-like element is located upstream of *flhDC* [27], which encodes the master operon of flagellar synthesis, motility, and chemotaxis [13]. Transcription and translation of *flhDC* is essential for the downstream expression of flagellar synthesis, motility, and chemotaxis genes [28]. Analyzing the sequences through breseq [26], we interpreted the sequencing data in the following manner: in presence of phage χ, the evolving bacterial populations were enriched for mutants where DNA immediately downstream of IS1 was deleted. The deletions began at the IS1 boundary and extended into *flhDC* and downstream genes, with lengths from 0.6 to 13.8 kb depending on the lineage. Such IS1-mediated adjacent deletions remove *flhDC*, resulting in a lack of flagellar regulon transcription (**Fig 2**C,D; n = 6 sequenced populations).

### Bacteria recover motility later in experimental evolution

Later in the evolution experiment (by passages 3 or 4), visible swim rings were observed on all Treatment-population plates (**Fig 3**B), indicating that motile, phage-resistant variants had emerged. Interestingly, lineage T10 did not show such growth at passage 10, possibly due to phage-driven extinction of host bacteria (**Fig S1 in S1 Appendix**). To determine the genetic changes in these Treatment populations, we obtained consensus whole-genome sequences from each Treatment lineage at passages 5 and 10, and compared these data to the ancestral (reference) genome [26]. Genes with polymorphic sites which were over 10% different than the ancestor sequence are listed in **Table S1** in S1 Appendix (passage 5) and **Table S2 in S1 Appendix** (passage 10). Notably, several of these genes are involved in chemotaxis, the synthesis of flagellar machinery, or the regulation of flagellar motility (7/41 at passage 5, 13/70 at passage 10). Mutations that occurred at frequencies >70% are shown in **Fig S2 in S1 Appendix**: 5/9 and 6/13 (respectively at passages 5 and 10) of the genes carrying these mutations were motility or chemotaxis-related. A large number of high-frequency mutations were found in *fliC*, the gene encoding the flagellin protein, which forms the flagellar filament monomer (**Fig 3**A, **Fig S2 in S1 Appendix**). FliC consists of conserved N- and C-terminal D0 and D1 domains and variable central domains D2 and D3 (**Fig S3** in S1 Appendix). A substantially smaller number of mutations were observed at >10% frequency in the Control populations, none of which were in FliC (**Table S3** in S1 Appendix).

**Fig 3.**
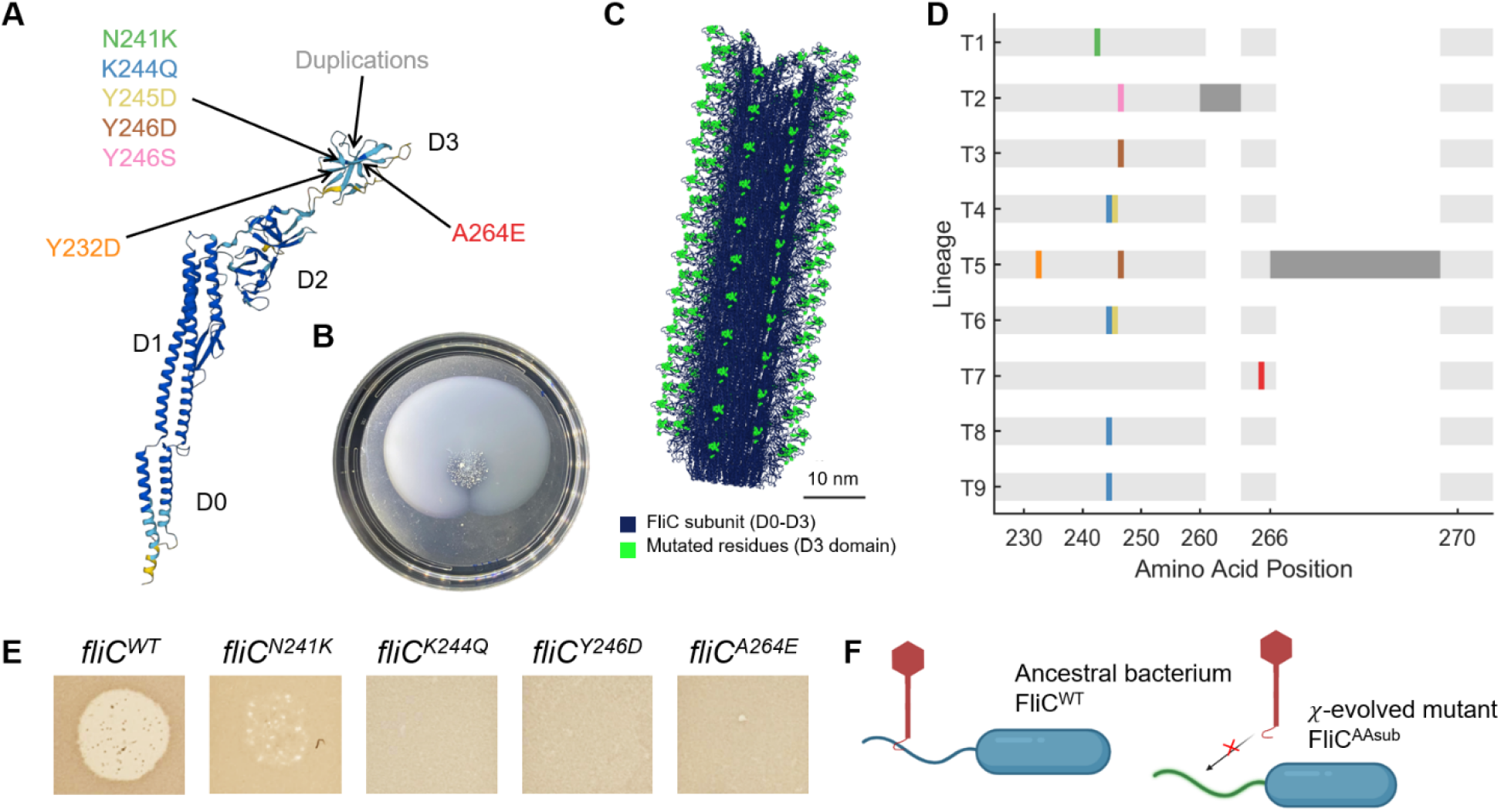
Motile, χ-resistant bacteria feature mutations in the flagellar filament. **(A)** Structure of *E. coli* FliC (UniProt P04949) is shown with domains D0, D1, D2, and D3. All observed *fliC* mutations mapped to the D3 domain. A zoomed-in view of the D3 domain with these amino acid (AA) residues is included in **Fig S3** in S1 Appendix. **(B)** All treatment populations showed visible swim rings after passage 4. **(C)** The mutated residues decorate the outer surface of the assembled flagellar filament. Composite structural model of the *E. coli* flagellar filament; dark blue: FliC subunits; bright green: mutated D3 positions; scale bar: 10 nm. See Materials and Methods for details of model generation. **(D)** Amino acid substitutions in domain D3 of FliC are indicated for each endpoint lineage (colored lines). Dark grey corresponds to AA sequences that were duplicated (7 AA duplication starting at T260 in T2; 19 AA duplication starting at A266 in T5). **(E)** Phage χ causes characteristic clearing on lawns of the ancestor, while cells carrying any of the single AA substitutions, N241K, K244Q, Y246D, and A264E in FliC are resistant to χ. **(F)** These results suggest a mechanism in which single amino acid substitutions in the native *fliC* locus of MG1655 (*fliC^AAsub^*) prevent χ from binding to the flagellum, allowing cells to evade infection. Panel F was created in BioRender. Turner, P. (2026) https://BioRender.com/51hbpmz.

We isolated multiple representative mutants (clones) from each endpoint (passage 10) Treatment lineage (at least two and up to seven clones isolated from each lineage) and performed Sanger sequencing of the *fliC* locus. All clones isolated from a single lineage carried identical mutations in *fliC*, so we report the consensus for each Treatment lineage (**Fig 3**D). Control lineages did not carry any mutations in *fliC*. Most mutations in the Treatment lineages were single or dual amino acid (AA) substitutions localized to the D3 domain of the flagellin monomer, which remains exposed on the outer surface of the flagellar filament after polymerization [29] (**Fig 3**C). Given the appreciable impact of point mutations on flagellar function [30,31], we hypothesized that a single AA substitution might confer resistance to phage χ. To test this idea, we constructed multiple strains via allelic exchange carrying our observed single AA substitution variants: FliC^A264E^, FliC^K244Q^, FliC^N241K^, and FliC^Y246D^ (**Fig 3**D). When we spotted dilutions of a 10^10^ PFU/mL phage χ stock onto lawns of these strains, we observed that clear plaques formed only on the ancestral wild-type (**Fig 3**E), indicating that cells expressing the AA substitutions escaped infection. We note that small individual plaques were also observed only at the highest χ titer (10^10^ PFU/mL) for FliC^N241K^. Images of full plates with 10-fold dilution series (10^10^–10^1^) are shown in **Fig S4** in S1 Appendix. We performed classic adsorption assays with two of these mutants, which revealed that χ does not adsorb to these strains (**Fig S5** in S1 Appendix). These results suggest that a single AA substitution in the flagellin’s D3 domain obstructs the binding of χ, rendering the bacteria resistant to the phage (**Fig 3**F).

We note that IS1-adjacent deletions were not reversed in later passages. They were likely outcompeted by motile, χ-resistant cells that never carried an IS1-mediated <u>deletion</u> (see Supplementary note in S1 Appendix).

### Phage **X**-resistance trades off and trades up with bacterial motility in soft agar

Two traits under selection can evolve via trade-offs (one trait improves at the other’s expense), trade-ups (both traits improve together under the imposed selection regime, without implying a context-independent fitness benefit), or may have neither relationship [32,33]. Since χ exploits the flagellar motility apparatus for binding, and motile and resistant cells carried mutations in the flagellar filament, we examined whether motility trades up or trades off with the evolution of χ-resistance, using clones isolated from each of the nine motile endpoint Treatment lineages and the four Control lineages. Swim-ring diameters after 15 hours were compared to those of the ancestor and Controls, which were serially passaged on swim plates in the absence of phage selection. For each Treatment and Control lineage, we experimented with each clone (isolated from passage 10 of each lineage) in duplicate. When tested individually, Control lineages did not show statistically significant differences in swim-ring diameter relative to the ancestor (which we attribute to low statistical power owing to small sample size); however, when pooled, the Controls collectively showed significantly larger swim rings than the ancestor (P < 10^-5^), consistent with soft-agar passaging itself driving modest improvements in motility performance even in the absence of phage selection [34,35]. Contrarily, most χ-resistant Treatment lineages (T1-T6) showed swim-ring diameters less than that of the ancestor, indicating an evolutionary trade-off: increased phage-resistance at the expense of decreased motility in soft agar (**Fig 4**A). Lineage T7 showed a swim-ring diameter comparable to that for the ancestor and the evolved Control strains. However, evolved motile lineages T8 and T9 exhibited significantly larger swim rings than the ancestor (**Fig 4**A, **Fig S6**A in S1 Appendix). When Treatment lineages were compared against the pooled Control dataset, T9 showed significantly greater swim-ring diameters than the pooled Controls, demonstrating a trade-up: increased phage-resistance alongside increased motility on soft agar. Lineages T7 and T8 recovered motility performance comparable to the pooled Controls despite continuous phage selection (**Fig S6**B in S1 Appendix).

**Fig 4.**
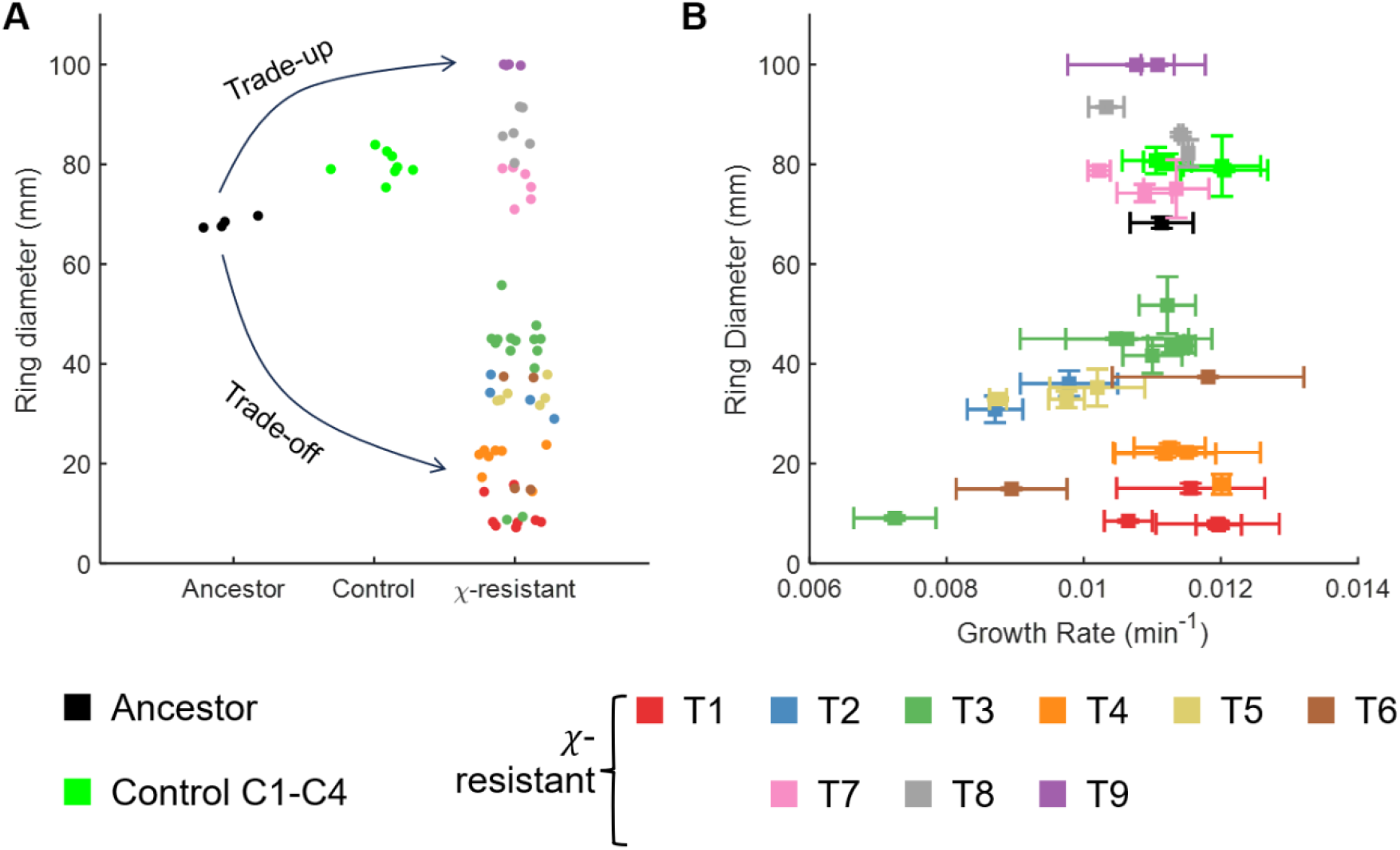
Resistance to phage χ trades off as well as trades up with motility. **(A)** Swim-ring diameters observed for the ancestor (black), strains isolated from evolved Controls (phage free; bright green), and motile χ-resistant Treatment (phage selection) lineages (color-coded according to Treatment population). Swim-plate assays were conducted in absence of phage χ. Most treatment populations showed evidence of either a motility trade-off with increased phage-resistance, or a motility trade-up with greater phage-resistance (trend indicated by arrows). **(B)** Growth rate of each bacterial strain was measured in well-mixed liquid (on a microplate reader) and plotted against the swim-ring diameter. Growth rate is reported as r_g_ in the exponential growth model *B(t) = B_0_ exp(-r_g_t)* where bacteria starting from initial density *B_0_* grow to density *B(t)* at time *t*. These observations were uncorrelated (Pearson correlation coefficient, ρ = 0.22, P-value = 0.2). Data points are color-coded based on the Treatment lineage for isolation. For each lineage, at least two (e.g., T6, T9) and at most seven (T3) clones were isolated. All experiments were carried out in duplicate for each clone. Further statistical tests are reported in **Fig S6** and **Table S4** in S1 Appendix.

In an expanding ring of motile bacteria on soft-agar plates, growth (cell division), flagellar motility, and chemotaxis all influence the expansion speed [13,36–39]. Recent studies have shown that population growth rates greatly affect expansion speeds [13,34,40]. To test whether our observed differences in swim-ring diameter can be attributed to varying growth rates, we measured the growth rates of the isolated endpoint clones (passage 10) using an automated spectrophotometer (microplate reader) where light scattering provides a proxy for cell density. We note that these growth rate measurements were performed in liquid culture rather than on solid or semi-solid medium; while this approach allowed high-throughput, replicate-rich measurement of growth independent of motility and chemotaxis, we cannot exclude the possibility that relative growth rates among clones differ on soft agar compared to liquid medium. We plotted the swim-ring diameter of each strain or clone against its growth rate (**Fig 4**B). These measured quantities showed no significant correlation (Pearson correlation coefficient ρ = 0.22, P = 0.2), suggesting that differential growth rate is not the primary factor influencing swim-ring diameter of the isolated treatment clones. Instead, mutations in the flagellar filament (**Fig 3**) and other related genes (such as genes encoding components of the flagellar motor or the chemotaxis pathway which is an upstream controller of flagellar motility, **Fig S2 in S1 Appendix**) likely alter the chemotaxis-ability (measured as the ‘chemotactic coefficient’ [13,36–39]) of the evolved Treatment strains in nutrient soft agar [41].

### **X**-resistance alters single-cell distributions of motility traits

After ruling out a general growth effect, we hypothesized that differences in chemotactic efficiency and/or motility may account for the observed variations in swim-ring diameters. To test this idea, we quantified single-cell swimming behavior through phase-contrast microscopy and single-particle tracking. We performed these experiments for the ancestral and evolved strains with a single representative clone from each end-point (passage 10) Treatment and Control lineage, as the flagellar mutations observed across all strains in each Treatment lineage were identical (**Fig 3**). We confirmed for each representative clone that motile cells remained resistant to χ (**Fig S7 in S1 Appendix**). We observed that a majority of cells in some of these clonal populations were non-motile and hence calculated the fraction of motile cells for each of the experimental lineages (**Fig 5**A). We observed that clones isolated from Treatment populations with reduced swim-ring diameters (lineages T1-T6) had significantly lower motile fractions than the ancestor. In contrast, the motile fraction in the remaining Treatment lineages (T7-T9) was comparable to the ancestor and the Control lineages (**Fig 5**A).

**Fig 5.**
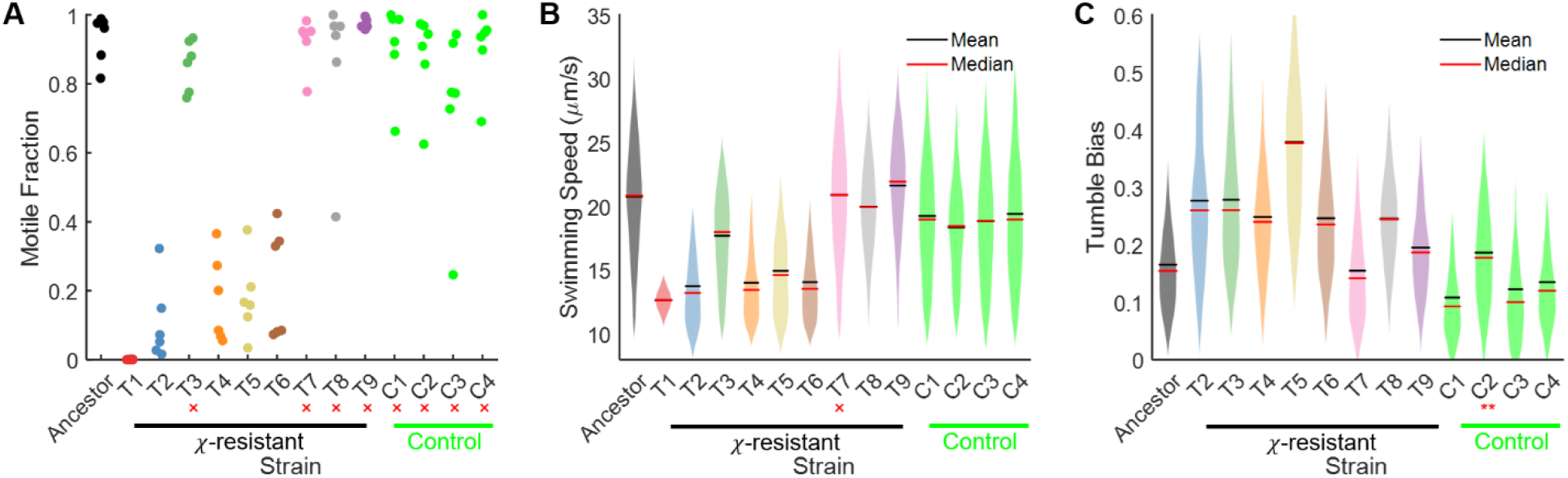
Motile fraction, swimming speed, and tumble bias of evolved strains often differ from traits of the ancestor. Single-cell motility was recorded with phase-contrast microscopy and analyzed using automated algorithms. **(A)** Motile fraction was calculated for a representative strain from each evolved lineage. **(B)** Distributions of single-cell swimming speeds are shown for each strain. **(C)** Tumble bias distributions of single cells are indicated for each strain (except χ-resistant strain T1, which was essentially non-motile). ANOVA yielded P << 10^-6^ for each dataset. Post-hoc Dunnett’s multiple comparisons tests were performed to compare each evolved strain with the ancestor. Strains marked with a red cross (×) showed no significant difference from the ancestor, while all the other observations differed significantly from the ancestor: P < 10^-4^ for all observations (not indicated) except tumble bias distribution for strain C2 which had P < 0.01, indicated with **). Exact P-value for each comparison is reported in **Table S4** in S1 Appendix.

We also compared the Treatment datasets with pooled Control datasets (**Fig S6**C-E in S1 Appendix), which revealed that motile fractions for T3 and T7-T9 were comparable to the pooled Controls (**Fig S6**C in S1 Appendix). Mean (or median) swimming speeds of T1-T6 cells were lower than pooled Controls, while those of T7-T9 were higher (**Fig S6**D in S1 Appendix). Each Treatment lineage demonstrated significantly elevated tumble bias as compared to pooled Controls (**Fig S6**E in S1 Appendix). Finally, we also compared the single-cell and population motility of the single AA substitution mutants (constructed in the ancestor background) to those for WT; the data are reported in **Fig S8** in S1 Appendix.

### **X**-resistant mutants have altered chemotaxis kinase activity

We expected the flagellar mutations to modulate motile fraction and swimming speeds, because change in flagellar structure (through AA substitutions characterized in **Fig 3**) may modulate the function. However, the change in tumble bias was somewhat surprising, as it is reflective of the chemotaxis response in wild-type cells. Albeit, it cannot be ruled out that mutant flagellar filaments cause the formation of an altered flagellar bundle, which could affect bias behavior. To evaluate the chemotaxis response of evolved isolates independent of flagellar filament function, we used a FRET-based approach to measure chemotaxis kinase activity in the ancestor strain and the evolved isolates [22,23]. In this assay, the addition and removal of a saturating concentration of the non-metabolizable chemoattractant α-methyl-aspartate (MeAsp) allows the determination of the chemotaxis kinase activity. An example trace of the kinase activity is shown in **Fig 6A**: upon addition (removal) of the attractant, the kinase activity goes down (up) and ‘adapts’ back to the basal value over time [22,23]. From the trace, we calculated three parameters of the kinase response: basal activity (a_O_), rate of adaptation (r), and precision in adaptation (a_adap_/a_O_), since adaptation to stimulus addition is typically imperfect for large ligand concentrations [42]. We measured chemotaxis kinase activity for the ancestor and three representative evolved bacterial strains: one clone isolated from an evolved trade-off lineage, T5, for which resistance to χ coincided with decreased motility, and two clones isolated from evolved trade-up lineages, T8 and T9, for which χ-resistance coincided with increased motility (relative to the ancestor). Parameters of the kinase responses for each strain are indicated in **Fig 6B,C,D**. The basal activity of the chemotaxis kinase appeared to be elevated in all evolved strains as compared to the ancestor, and this elevation was statistically significant for T5, with low variance across the three replicates (**Fig 6B**). Trade-up strains T8 and T9 showed significantly faster kinase adaptation as compared to the ancestor, while trade-off strain T5 was comparable to the ancestor (**Fig 6C**). The adaptation was more precise in the trade-up strain T8 (and showed an upward trend in T9), while the trade-off strain T5 adapted less precisely as compared to the ancestor (**Fig 6D**). Overall, we observed that the motility-trade-up strains T8 and T9 evolved to adapt faster and more precisely, while the trade-off strain T5 did not.

**Fig 6.**
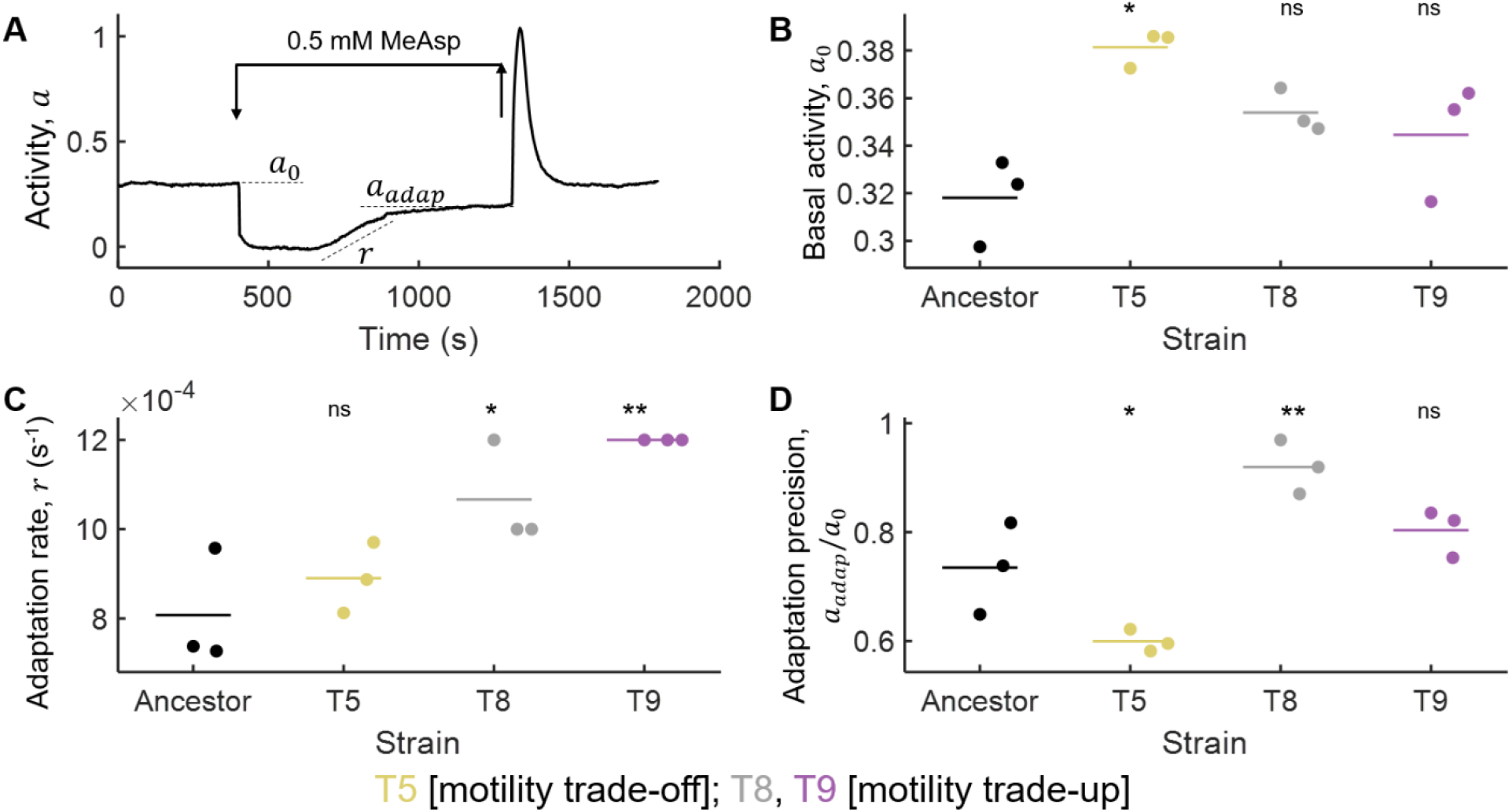
Evolved strains respond differently to chemotactic stimuli. **(A)** An example trace of the kinase activity (ratiometric FRET between CheZ-YFP and CheY-RFP) is shown. The arrows indicate the time when a saturating concentration of chemoattractant methyl-aspartate (0.5 mM) was flowed into and out of the sample chamber. Basal activity a_0_, adapted value of activity a_adap_, and adaptation rate r are indicated with dashed lines (r is calculated as the slope of the line, whereas a_0_ and a_adap_ are Y-values). Each trace is an output from observations of hundreds of cells within the microscopic focal view. Basal activity a_0_ **(B)**, adaptation rate r **(C)**, and adaptation precision calculated as the ratio of the adapted and basal values of kinase activity, a_adap_/a_0_ **(D)** for three independent replicated experiments are indicated as dots for each strain. Mean values are indicated as dashes. For each parameter in B-D, ANOVA yielded P < 0.01 for each dataset (P = 0.01, 0.0041, and 6.38 × 10^-4^ respectively, for B, C, and D). Post-hoc Dunnett’s multiple comparisons tests were performed to compare each representative lineage strain with the ancestor (* P < 0.05, ** P < 0.01, ^ns^ no significant difference from the Ancestor group). Exact P-value for each comparison is reported in Table S4 in S1 Appendix.

We also performed FRET experiments on two Control lineages, indicated in **Fig S11** in S1 Appendix. Control lineages do not deviate substantially from the ancestor in their mean chemotaxis signaling parameters (though individual lineages showed significant deviation in certain parameters), but show markedly higher variability across replicates compared to both the ancestor and Treatment lineages. We attribute this to the increase in cell-to-cell phenotypic variability that arises from growth in soft agar without directional selection, as recently reported in an independent study [43]. Because these are population-average FRET measurements, the elevated replicate-to-replicate variability in Controls reflects this underlying single-cell heterogeneity. This contrast is itself informative: strong χ-phage selection produced consistent and significant modulation of chemotaxis signaling parameters across Treatment lineages (**Fig 6**), whereas passaging without phage selection yielded no directional shift but increased variance (**Fig S11** in S1 Appendix). Including Control lineages in the Treatment-vs-ancestor adaptation-rate comparison abolishes some of the significant differences reported above, which reflects this added Control variance rather than a loss of the Treatment-specific effect (**Fig S11** in S1 Appendix).

## Discussion

Flagellotropic phages are commonly found in natural samples and may have existed alongside bacteria for billions of years of evolution [5,6,16,44–49]. Motile bacteria may regularly encounter flagellotropic phages, yet the evolutionary mechanisms of bacterial resistance to these phages are rarely studied [14]. We examined the genetic and phenotypic consequences for bacteria when they evolve in microcosms that allow cell motility and contain flagellotropic phage predators, using the model system bacterium *E. coli* and phage χ. Instead of using well-mixed environments (e.g., shaking culture-flasks) typical of many phage-bacteria coevolution experiments, we evolved bacteria in soft-agar swim plates to maintain spatial structure and preserve motility as a selectable trait. Critically, phage χ was uniformly pre-embedded throughout the agar at a concentration sufficient to suppress swim-ring expansion, eliminating spatial escape as an adaptive strategy and ensuring that all migrating cells encountered phage with high probability across the entire plate. This design differs from recent studies in which phage and bacteria were co-inoculated at a fixed point, allowing motile cells to physically outrun the phage front [50–52]—this is a strategy that selects for motility as a means of spatial escape rather than forcing evolutionary adaptation of the phage receptor itself. Our approach thus isolated a specific and underexplored question: how does sustained phage predation acting directly on the flagellum reshape the motility and chemotaxis phenotype through evolutionary adaptation? Beyond documenting that bacteria evolve χ-resistance through flagellin modification — a finding consistent with the broader principle of receptor evolution under phage predation — we show that phage selection simultaneously reshapes distributions of single-cell swimming speed and tumble bias, and chemotaxis signaling dynamics, revealing consequences for the entire motility machinery that extend well beyond the receptor itself.

Phage χ is known to infect both *E. coli* and *Salmonella enterica* [20,53], which possess a similar motility and chemotaxis machinery [18]. In *E. coli*, results from our evolution experiment show that all mutations that disrupt χ infection were in the outermost D3 domain of *E. coli* flagellin (**Fig 3**A, notably, no mutations to *fliC* were observed in the Control group of evolved lineages). A targeted domain deletion and swapping approach in *S. enterica* revealed that the N- and C-terminal D2 domains are the major determinants of phage susceptibility and that the D3 domain plays a less-crucial role [54]. These two results appear to be in some disagreement. Interestingly, we found no mutations in efflux pump genes in any Treatment lineage, and a Δ*tolC* mutant remained susceptible to χ (**Fig S9** in S1 Appendix), suggesting that unlike in *Salmonella enterica* serovar Typhimurium strain 14028 [53], TolC is not required for χ infection in *E. coli*.

In addition to the mutations in flagellar motor and chemotaxis machinery, some of our populations also acquired mutations in genes involved in the synthesis or regulation of outer-surface structures such as lipopolysaccharides (LPS) and capsules (**Fig 3**, **Table S1, Table S2 in S1 Appendix**). LPS-related mutations were also observed in earlier studies that imposed phage χ selection pressure on *E. coli* or *Salmonella* [55,56]. Flagellotropic phage 7-7-1 uses LPS as the secondary receptor to infect its host *Agrobacterium* sp. [45]. Model coliphage T4 also uses LPS as a co-receptor [57,58]. On the other hand, several mutations in LPS biosynthesis and Rcs pathways are known to have pleiotropic effects in reducing motility [59,60]. Our results suggest two possibilities: either that phage χ may use LPS in some capacity during the cell adsorption and genome injection processes, or, more likely, that reduced motility in the LPS biosynthesis and Rcs mutants may provide limited resistance against χ.

In the current model of χ-attachment to the flagellum, CCW rotation of the flagellum drives the phage towards the cell body [20]. In this model, clockwise rotation of the flagellum would propel the phage away from the cell. Our kinase activity measurements revealed higher mean kinase activity in the evolved lineages (**Fig 6**B: statistically significant difference in mean a_O_ for trade-off strain T5, with upward trends for T8 and T9). Higher kinase activity increases phosphorylation of CheY, where CheY-P promotes clockwise rotation of flagellar motors (**Fig** **1**A). Additionally, our single-cell tracking experiments showed that tumble bias increased in most evolved, χ-resistant lineages. This increased tumbling may result from either a higher fraction of clockwise (tumble-inducing) flagellar rotation or altered bundle stability due to AA substitutions in FliC. We do not believe our results indicate adaptive changes in tumbling specifically to resist χ attack; rather, we propose that AA substitutions in FliC confer resistance to χ by blocking adsorption (**Fig S5** in S1 Appendix), while elevated tumble bias may be a secondary phenotype that reduces the probability of χ reaching the cell body. More broadly, we acknowledge that the mechanistic link between the observed single-cell motility phenotypes and phage resistance remains to be established. The flagellin mutations identified in **Fig 3** are sufficient to confer χ-resistance, and the single-cell data document the collateral phenotypic consequences of those mutations on swimming behavior. Whether the specific changes in swimming speed or tumble bias are causally adaptive, pleiotropic byproducts of flagellin structural changes, or incidental outcomes of evolution in soft agar is an open question we intend to pursue as a follow-up to this work.

Recent work has shown remodeling in motility as well as the chemotaxis network and specifically kinase CheA activity, during evolution of motile *E. coli* on swim-plates [39,61]. The Treatment strain T5 (that evolved a motility trade-off with χ-resistance) exhibited elevated basal CheA activity and reduced adaptation precision as compared to the ancestor. In contrast, the trade-up strains T8 and T9, which improved both motility and χ-resistance, likely recovered their performance by evolving faster and more precise adaptation (**Fig 6**). The fact that the trade-up strains adapt faster than the ancestor likely explains how these strains expand faster without having a considerably higher swimming speed or growth rate [42]. We note that differences in CheA kinase activity across lineages could, in principle, reflect altered expression of flagellar or chemotaxis genes rather than direct adaptive remodeling. However, the observation that trade-up lineages (T8, T9) evolved faster and more precise kinase adaptation (**Fig 6**), which is a phenotype mechanistically consistent with their expanded swim-ring diameters (**Fig 4**), suggests these are not incidental expression-level differences. The AA substitutions in FliC are unlikely to be responsible for the changes in CheA response; rather, other mutations within chemotaxis and motility genes (**Table S1**, **Table S2 in S1 Appendix**) likely modulated the kinase response. Future experiments using recombinant genetics could resolve which specific mutation(s) were responsible for the observed changes in kinase activity. We believe this is the first evidence of evolutionary pressure from a phage remodeling the dynamics of regulatory pathways (chemotaxis signaling in this case) upstream of the phage receptor (flagella in this case).

The century-old approach of using phages as therapeutics is experiencing resurged interest due to the alarming rise in antibiotic-resistant bacterial infections [4]. Evolutionary trade-offs can be leveraged in phage therapy, such that target bacteria are killed while therapeutic phages select for mutants that resist phage attack alongside phenotypic changes that reduce pathogenicity [1]. However, trade-ups may also occur, potentially making the target bacteria resist phage infection while becoming more pathogenic [32]. Because motility is a fitness or virulence factor in many bacterial species [15], flagellotropic phages offer exciting therapeutic options [5]. Our results demonstrate that *E. coli* cells exposed to phage χ first evolve resistance by forfeiting motility, implying a potentially advantageous fitness and/or virulence trade-off. In a clinical setting, the remaining non-motile bacteria would be less capable of host colonization and could be more readily eliminated by immune defenses or co-therapy with antibiotics. Thus, the early stages of ‘flagellotropic-phage-steered’ bacterial evolution are relevant in therapy development, while caution is needed when considering later stages of evolution due to the possibilities for trade-ups.

According to later stages of our experiments, persistent phage selection may favor χ-resistant cells that preserve motility. In our model system of *E. coli* and χ phage, this balance was achieved through mutations in the flagellar filament that preclude χ-binding without disabling swimming. Thus, our results suggest that the evolutionary response of bacteria to flagellotropic phages is not straightforward and may depend on the duration of phage selection in addition to random mutational variation across populations.

## Materials and Methods

### Bacteria and phage strains

MG1655 was used as the wild-type (WT) *E. coli* ancestor strain, which was obtained from J. Wertz at the Coli Genetic Stock Center (CGSC) at Yale University, now independently operating as *E. coli* Genetic Resource Center (https://ecgrc.net/). *Salmonella*-infecting phage χ was obtained from Kelly Hughes at the University of Utah and grown on the permissive (Δ*recA*) strain BW25113 to change its methylation signature to *E. coli*.

### Mutant construction in *E. coli* MG1655

Point mutations in *fliC* were generated in vitro by overlap extension PCR. Constructs of approximately 1,000 bp in length containing the mutation were cloned into the suicide vector pWM91. The diamino pimelic acid (DAP) auxotrophic *E. coli* strain B2155 was transformed with the pWM91 derivatives. Recipient and donor strains were grown to an OD_600_ of 0.8 to 1.0 in LB or LB containing 100 μg/mL ampicillin (Amp) and 60 mM DAP, respectively. The donor strain was harvested by centrifugation for 4 minutes at 8,000 rpm, gently suspended in LB/DAP, and washed twice with LB/DAP. Recipient and donor strains were mixed in a 1:1 ratio, 100 μL of the mixture (mating mix) was pipetted on a cellulose acetate filter, which was placed on an LB/DAP plate, and incubated overnight at 37°C. The mating mix on the filter was suspended in LB and washed once with LB+Amp. Serial dilutions of the mating mix were plated on LB+Amp plates, and single colonies were streaked on selection plates. 10 mL of LB was inoculated with individual colonies and grown at 37°C with shaking. Cells were harvested by centrifugation at 8,000 rpm for 4 minutes in a tabletop centrifuge, the supernatant was aspirated, and the cell pellet was washed with LB_0_ (LB lacking NaCl). Serial dilutions were plated on LB_0_ containing 10% sucrose and incubated at room temperature for two days. Colonies were patched on LB plates with and without ampicillin. Ampicillin-sensitive colonies were screened by PCR and Sanger sequencing, and lack of growth on LB+Amp plates was confirmed via T-streak.

### Culture conditions

Tryptone Broth (TB; 10 g tryptone, 5 g NaCl per L) was used for liquid cultures, 1.5% agar plates, 0.5% top agar for the double-layer method of phage propagation and 0.3% swim agar for swim plate assays. Overnight cultures were initiated from colonies grown on solid (1.5%) TB agar and placed into TB liquid medium, incubated with shaking (150 RPM) at 30°C. Bacterial stocks were stored in 25% glycerol at −80°C. High-titer stocks (lysates) of phages were grown by mixing a virus with its wild-type host bacteria in liquid broth and incubating for 12–24 hours as described above to allow phage population growth, followed by centrifugation and filtration (0.22 μm) to remove bacteria and obtain a cell-free lysate. Phages were enumerated (plaque-forming units [PFU] per mL) using the standard double-layer agar method where viruses form visible plaques on confluent lawns of wild-type host bacteria within 0.5% top agar, overlayed on 1.5% agar in the bottom layer.

### Evolution Experiment

Treatment lineages of *E. coli* were passaged as follows. Soft-agar swim plates (Tryptone Broth [TB] with 0.3% agar) were embedded with a high concentration of phage χ to ensure that bacteria frequently encounter phages while swimming: 15 μL of 10^11^ PFU/mL χ were added to 10 mL of agar in each plate. This concentration of phages was determined by carrying out a preliminary experiment to test the lowest phage concentration that suppressed bacterial swim ring formation for 24 h. For the evolution experiment, a phage titer 10-fold greater than the threshold was used. 5 μL of overnight culture of MG1655 (the ancestral strain) grown in TB was inoculated in the center of the plates followed by incubation for 24–48 h at 30°C.

After each passage (until swim rings appeared or 48 hours elapsed), plates were processed as follows: the contents of the plate were pooled into a tube with 10 mL TB and homogenized by vortexing. Next, 10 μL of the mixture (effectively 5 μL of the previous plate’s contents, 1:2,000 evolutionary bottleneck) was inoculated into a fresh swim plate containing phage χ at the same concentration as indicated above. Plates were incubated at 30°C and a frozen stock in 25% glycerol was made from the remaining mixture. This process was repeated for 10 passages, with 10 independent replicates all initiated from the same MG1655 ancestor genotype (**Fig 1**C). Each new passage occurred after 24 hours if a motile ring of bacteria was observed. If no motile ring was present (which was the case for the first 2-3 passages), incubation continued until 48 hours. This method allowed us to determine if the χ-resistant bacteria growing on the plate remained motile. Four Control populations of *E. coli* were passaged identically, in the absence of phage.

### Isolation of bacteria from evolution experiment

At least two individual isolates (clones) were obtained from each Treatment and Control lineage at passage 10. The frozen stock was streaked on 1.5% TB agar plates to obtain isolated colonies. Individual isolated colonies were picked and re-streaked onto fresh plates. Individual colonies from the second (doubly-isolated) plate were used for experiments with the isolates (**Fig 3**, **Fig 4**).

### Sequencing and mutant calling analysis

For whole genome sequencing of each Treatment lineage at passages 5 and 10, 200 μL of frozen stock were placed in 5 mL TB and incubated overnight at 30°C. After incubation, 1 mL of each culture was pelleted by centrifugation and sent to SeqCenter, where genomic DNA was extracted and libraries were prepared for Illumina sequencing (paired-end reads of 151 bp length collected to a final coverage of ∼100-fold across the reference genome). Mutant calling analysis was performed using breseq with polymorphism mode [26]. Read depth was computed from the breseq alignments with pysam, and junctions joining IS1 to adjacent sequence were taken from breseq’s junction evidence. Partial deletions are not reported as mutations by breseq in polymorphism mode.

### Rendering of mutated residues onto flagellar filament

The outer domains of *E. coli* FliC are unresolved in its only experimental structure (PDB 8CXM models residues 3–177 and 407–498 only), so a composite model was built. Core coordinates of domains D0 and D1 were taken from 8CXM. The D2/D3 fold was taken from the AlphaFold Database model of P04949 (AF-P04949-F1) and positioned by superposing its D2 domain onto that of the *Salmonella* full-length filament PDB 3A5X (1.9 Å RMSD over 527 atoms), with the two placements blended across the hinge. This filament from *Salmonella* was used as the outer-domain reference because *E. coli* K-12 filaments are canonical and non-sheathed, unlike the fully resolved structures of *E. coli* outer-domain-sheath filaments [62]. The subunit was propagated by helical parameters fitted to 8CXM (−65.40° twist, −4.83 Å rise per subunit) to give a 64 nm filament of 121 subunits, 23.3 nm in diameter, rendered in PyMOL 3.1.0.

### Swim ring diameter measurements

Swim ring diameters were determined using a previously published protocol with some modifications [53]. Briefly, 2.5–5 µL of stationary phase culture was spotted at the center of freshly prepared TB soft agar (0.3%). Plates were incubated at 30°C over 8–13 hours and swim ring diameters were measured (inoculum volume and incubation period varied across—but not within—experimental datasets).

### Measurement of phage adsorption

Overnight cultures were grown at 30°C in TB medium, diluted 1:100 in fresh TB, and re-grown until reaching OD_600_ values between 0.7 and 0.8, and phage was added at a final titer of ∼1×10^6^ PFU/mL to all cultures. Cells were incubated with phage for 10 minutes at 30°C while shaking at 226 RPM before being collected and centrifuged at 21,130 RCF on a tabletop centrifuge for 3 minutes. Phage-containing supernatants were titered via plaque assay.

### Spot assays

Spot assays were performed by combining motile, mid-log-phase cells with molten TB supplemented with 0.5% agar and overlaying the mixture onto LB plates. Overlayed agar was allowed to solidify before a series of phage serial dilutions was spotted in 2.5 µL droplets on the overlay. Plates were incubated upright overnight at 30°C.

### Bacterial growth rate measurements

Isolates were grown overnight in TB medium at 30°C, then diluted 1:200 into 200 μL fresh TB. Each isolate was assayed in triplicate with their positions on the 96-well plate randomized using the *PlateDesigner* application [63]. Cultures were incubated at 30°C with shaking, and optical density at 600 nm (OD_600_) was monitored at 10-min intervals for 24 h by a BioTek Synergy H1 microplate reader. The growth rate *r_g_* was calculated by fitting the exponential growth model *B(t) = B_O_ exp(-r_g_t)*, where bacteria starting from initial density *B_O_* grow to density *B(t)* at time *t*. Thus, doubling time *T = ln2/rg*. Taking OD_600_ as a proxy for bacterial density *B*, the growth rate *r* was extracted from the linear region of log(OD_600_) versus time for each strain.

### Single-cell tracking experiments

Overnight cultures were grown at 30°C in TB medium, diluted 1:100 in fresh TB, and re-grown for 4 hours at 30°C, reaching the exponential phase of growth (OD_600_ ∼ 0.4-0.5). Exponential-phase cells were diluted into Motility Buffer (MB: 0.01 M potassium phosphate, 0.067 M NaCl, 0.1 mM EDTA, 1 μM methionine, 10 mM lactic acid, pH 7.0) to an OD_600_ ∼ 10^-4^ and allowed to equilibrate (chemotactically adapt to MB) at room temperature for 20 min before experiments were performed. Cells were then introduced to an imaging chamber made by affixing #1.5 coverslips to glass slides via double-sided sticky tape. Both glass slides and coverslips were freshly treated (on the same day) with vacuum gas plasma to make their surfaces hydrophilic, thereby preventing adhesion of cells to these surfaces. After introducing the sample, the chamber was sealed on both sides with VALAP (equal parts Vaseline, lanolin, and paraffin wax) to prevent fluid flow. Swimming cells were visualized at 30°C with a Nikon Ti-E inverted microscope using a CFI Plan Fluor 4×/0.13 NA objective and PhL ring for phase-contrast. Movies were recorded at 20 frames per second and analyzed in MATLAB using algorithms described previously [38,64,65].

### Calculation of motile fraction, swimming speed, and tumble bias

Distributions of the swimming speed featured a peak ∼ 5 μm/s. Control experiments with the non-motile MG1655 Δ*fliC* strain confirmed the peak to be due to non-motile cells (**Fig S10 in S1 Appendix**). Because this peak terminated at 8-10 μm/s, cells with speeds < 10 μm/s were classified as non-motile. Therefore, the motile fraction in each experiment was defined as the portion of trajectories exceeding the 10 μm/s threshold.

Swimming speed and tumble bias were calculated as described previously [38,64]. Briefly, a custom MATLAB algorithm was used to categorize trajectory segments as ‘run’ or ‘tumble’. Tumbles were primarily identified using angular speed: local minima and maxima were defined, and a tumble was identified if the total angular change over a local angular velocity peak exceeded a threshold of 4√*D_r_Δt* (rotational diffusion constant assumed as *D*_r_ = 0.1 *rad_2_/s*). The tumble duration was defined as the interval where the angular speed remained above 50% of the peak value, provided it did not drop below an exit threshold of 15 *rad/s*. Additionally, linear speed criteria were employed to identify tumbles. A speed-based tumble was called if the depth of a local speed minimum (Δv, relative to adjacent maxima) was at least 2.5 times the minimum speed. For these events, the tumble was defined as the period where the speed remained within 25% of the minimum value relative to the recovery. Finally, any single-frame gaps between detected tumbles were filled to account for temporal down-sampling. For each trajectory, the mean swimming speed was calculated as the average translational speed during non-tumble periods, and the tumble bias was calculated as the ratio of time spent tumbling to the total trajectory time.

### *In vivo* FRET microscopy

Cells were transformed with plasmid pSJAB106, which expresses CheZ–YFP and CheY–mRFP1 fusion proteins in tandem (constituting a FRET pair) under IPTG induction, and contains an ampicillin-resistance cassette for selection [23]. Cells were grown overnight at 30°C in TB, supplemented with 100 µg/mL ampicillin to ensure plasmid retention. Overnight cultures were then diluted 1:50 into 10 mL of fresh TB supplemented with 50 µM IPTG and 100 µg/mL ampicillin and then grown at 30°C until they reached an OD_600_ of 0.45-0.46. Cells were subsequently washed twice with MB^-^ (Motility Buffer described above, without NaCl) and resuspended in 2 mL MB^-^. The suspension was incubated at room temperature for 90 min to allow fluorophore maturation.

For FRET imaging, 60 µL of cell suspension were incubated on a coverslip coated with poly-L-lysine (Sigma) for 10 min to attach cells, then transferred to a flow cell under continuous flow of MB^-^ (400 µL/min) maintained by a syringe pump (PHD 2000, Harvard Apparatus). Solutions in MB^-^ were used to add and remove 500 µM α-methyl-aspartate (MeAsp; Sigma).

FRET imaging was performed on an inverted microscope (Eclipse Ti-E, Nikon) equipped with a 60× oil-immersion objective (CFI Apo TIRF, Nikon). YFP excitation took place every 2 s using a broad-spectrum LED (SOLA SE, Lumencor) with a pulse duration of 50 ms. Excitation light was passed through two excitation filters (59026x, Chroma; FF01-500/24-25, Semrock) and a dichroic mirror (FF520-Di02, Semrock). Emitted light was split into two channels using an image splitter (OptoSplit II, Cairn Research) equipped with a dichroic mirror (FF580-FDi01-25×36, Semrock) and two emission filters (FF01-542/27 and FF02-641/75, Semrock), projecting YFP and RFP signals side-by-side onto a single sCMOS camera (ORCA-Flash 4.0 V2, Hamamatsu). A dense monolayer of approximately 500 cells was imaged in each experiment. Images were binned 4×4 to maximize signal-to-noise ratio. All experiments were carried out at room temperature (∼22°C).

To correct for the drift in fluorescence intensities primarily due to fluorophore bleaching, the population FRET ratio (defined as the ratio of the raw emitted RFP and YFP signals) was fitted with a double exponential function. Subsequently, the FRET ratio was divided by the double exponential and normalized between zero (activity upon addition of a saturating dose of 500 μM attractant MeAsp) and one (activity upon attractant removal). Hence, the kinase activity was calculated as:

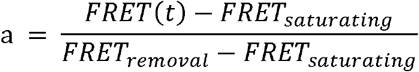

### Use of AI-assisted tools

Claude Opus 5 was used for the following purposes:

i. to construct the composite structural model described above and to generate the molecular renderings in **Fig 3**C and **Fig S3** in S1 Appendix, using open-source PyMOL 3.1.0. No image content was generated by AI; these figures are renderings of atomic coordinates. Model geometry was validated by the authors against the deposited structures.
ii. to assist with the flagellin sequence alignment analysis reported in **Fig S3**A, B in S1 Appendix; further details are provided in the figure legend.
iii. To assist in structuring the online data repository with systematic README files. Claude Sonnet 4.6 was used for language editing.

## Data Availability

The raw sequencing reads generated in this study have been deposited in the NCBI Sequence Read Archive (SRA) under BioProject PRJNA1544575. This BioProject will be published upon publication of the peer-reviewed manuscript.

The minimal dataset and accompanying code is available at Yale Dataverse via https://doi.org/10.60600/YU/KO5OXK. This repository will be published for the general audience upon publication of the peer-reviewed manuscript.

## Acknowledgements

We thank Kelly Hughes for gifting us a stock of phage χ, John Wertz for his help in generating the *fliC* knockout for this work, Ece Karatan for plasmid pWM91, Jun Zhu for *E. coli* B2155, and Paul Cohen for MG1655Δ*fliC*Δ*motAB*. We thank Turner Lab members including Noah Houpt, Kaitlyn Kortright, Catherine Hernandez, Michael Blazanin, Dallas Mould, Albert Vill, Helen Stone, and others as well as Jeremy Moore, Lam Vo, Jake Sumner, and other members of the Yale Quantitative Biology Institute for interesting discussions and valuable feedback about this work. We also thank Scharf lab members Madeline Harris and Eli Konvalinka for assisting with adsorption assays and swim ring diameter measurements.

## Supporting information

**S1 Appendix. Supplementary figures, tables, and note.** Contains Fig S1–S11, Table S1–S4, and the Supplementary note on IS1-mediated deletions.

**S1 Data. Additional details for Table S1 in S1 Appendix.** Genes with polymorphic sites over 10% different from the ancestor sequence at passage 5 in Treatment lineages (tab-separated values).

**S2 Data. Additional details for Table S2 in S1 Appendix.** Genes with polymorphic sites over 10% different from the ancestor sequence at passage 10 in Treatment lineages (tab-separated values).

**S3 Data. Additional details for Table S3 in S1 Appendix.** Genes with polymorphic sites over 10% different from the ancestor sequence after passage 10 in Control lineages (tab-separated values).

