## Supplementary material for "Flagellotropic bacteriophage selects for evolutionary remodeling of cell motility and chemotaxis in *Escherichia coli*": S1 Appendix

[Short title] Phage Selection Remodels Bacterial Motility

**Jyot D. Antani[1,2,3,✉], Austen Theroux[1], Fotios Avgidis[3,4], Jackson A. Wood[5], Birgit E. Scharf[5], Thierry Emonet[3,4,6], Paul E. Turner[1,2,3,7,✉]**

Contents: Fig S1–S11, Table S1–S4, Supplementary note: IS1-mediated deletions.

### Supplementary Figures

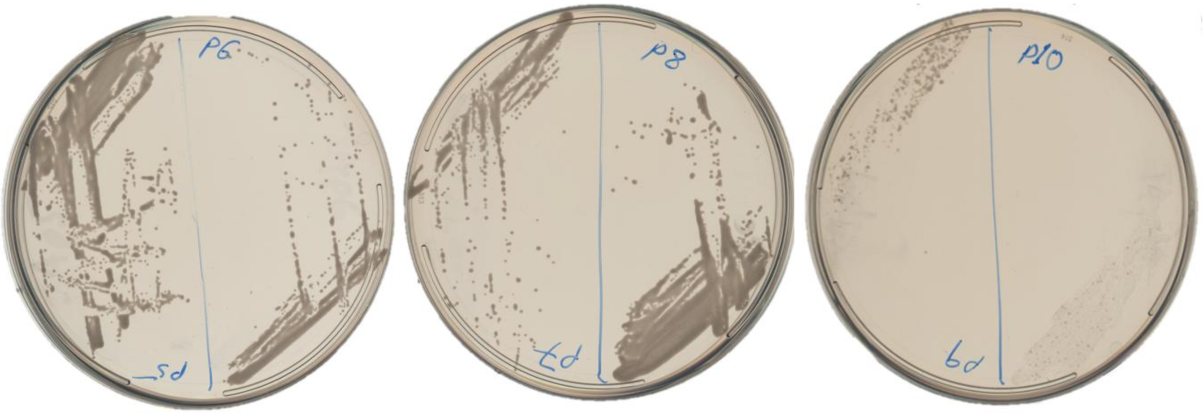


**Fig S1. Cells in lineage T10 went to extinction by passage 10.** Plates with colonies streaked out from frozen samples at the end of each passage starting at passage 5 are indicated.

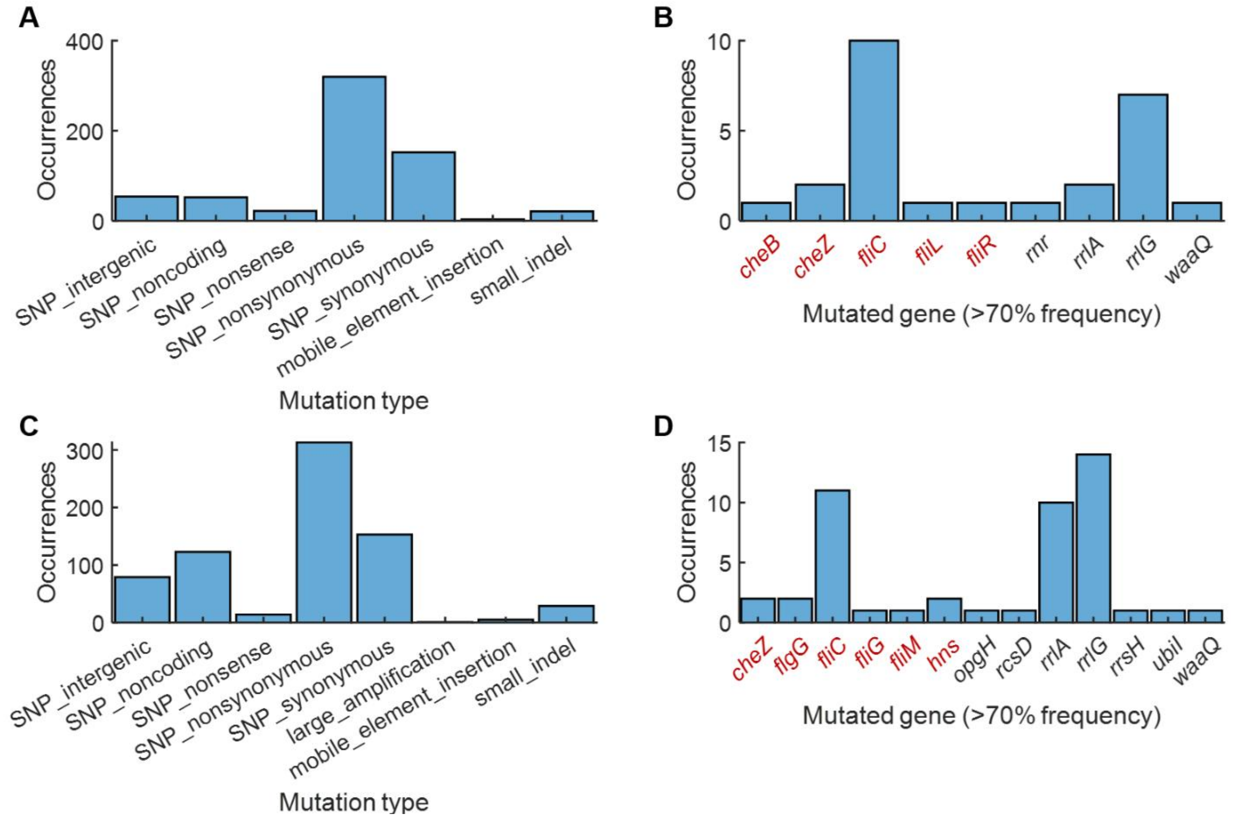


**Fig S2. Mutation frequencies in pooled genomes from all lineages (A, B) or all lineages except lineage T10 (C, D) are depicted. (A)** Number of occurrences of each type of mutation in passage 5. **(B)** Among mutations in passage 5 with frequency >70%, the number of occurrences of each gene are plotted. **(C)** Number of occurrences of each type of mutation in passage 10. **(D)** Among mutations in passage 10 with frequency >70%, the number of occurrences of each gene are plotted. Genes known to be related to chemotaxis and flagellar motility are indicated in dark red color.

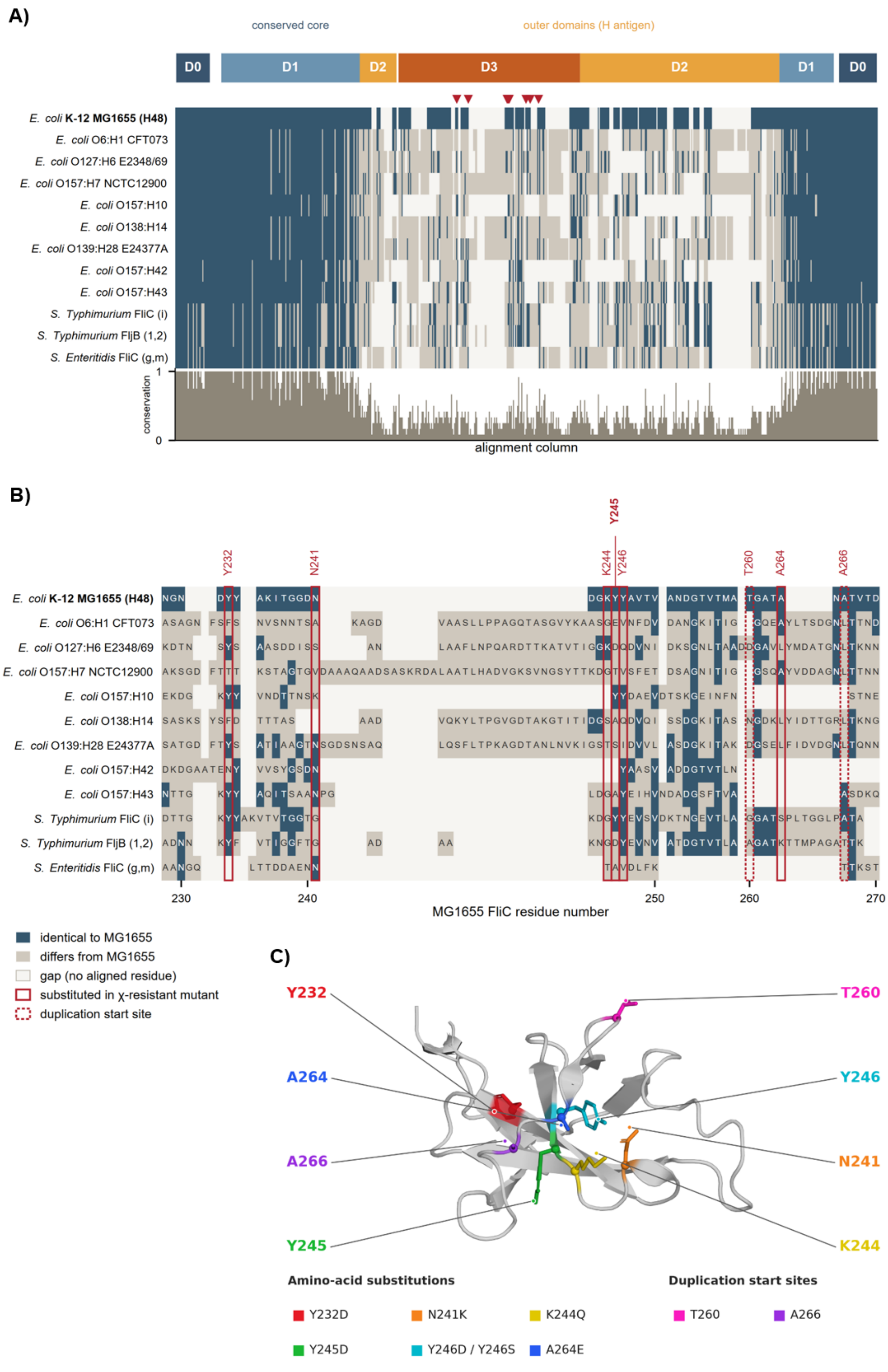
A)
conserved core
outer domains (H antigen)
D0
D1
D2
D3
D2
D1
D0
E. coli K-12 MG1655 (H48)
E. coli O6:H1 CFT073
E. coli O127:H6 E2348/69
E. coli O157:H7 NCTC12900
E. coli O157:H10
E. coli O138:H14
E. coli O139:H28 E24377A
E. coli O157:H42
E. coli O157:H43
S. Typhimurium FliC (i)
S. Typhimurium FljB (1,2)
S. Enteritidis FliC (g,m)
conservation
1
0
alignment column
B)
Y232
N241
K244
Y245
Y246
T260
A264
A266
230
240
250
260
270
MG1655 FliC residue number
identical to MG1655
differs from MG1655
gap (no aligned residue)
substituted in χ-resistant mutant
duplication start site
C)
Y232
T260
A264
Y246
A266
N241
Y245
K244
Amino-acid substitutions
Y232D
N241K
K244Q
Y245D
Y246D / Y246S
A264E
Duplication start sites
T260
A266

18

**Fig S3. The $\chi$-resistance mutations cluster in the hypervariable D3 domain of flagellin. A)** Domain architecture of FliC and a multiple-sequence alignment of twelve flagellins colored by identity to *E. coli* K-12 MG1655. *[Top]* the flagellin monomer's domain organization: the conserved inner core (D0, D1) and the surface-exposed outer domains (D2, D3) that carry the H-antigen. D1 and D2 are each discontinuous in sequence, contributing an N- and a C-terminal segment that flank D3. *[Middle]* each row is one flagellin: *E. coli* K-12 MG1655 (H48; the evolved ancestor), eight further *E. coli* H-antigen types, and *Salmonella* Typhimurium FliC and FljB and *S.* Enteritidis FliC (accession numbers along with code to reproduce these results are included in data repository). The flagellins were aligned with MAFFT L-INS-i and colored per residue as identical to MG1655 (dark blue), differing (tan), or gap (pale). Red triangles above the MG1655 row mark the eight D3 positions altered in $\chi$-resistant mutants. Domain boundaries are from the CATH structural assignment of the *Salmonella* flagellar filament (PDB 3A5X), transferred to MG1655 through the alignment (D3 = MG1655 residues 196–282). *[Bottom]* per-column conservation (fraction of the twelve sequences identical to MG1655). **B)** Residue-level view of the outer D3 domain (MG1655 FliC 228–270), colored as in **(A)**. The six substituted positions are boxed with solid lines and the two in-frame–duplication start sites with dashed lines. **C)** A zoomed-in view of the FliC D3 domain is shown with key AA residues highlighted in color. Because the outer domains of *E. coli* flagellin are unmodelled in the only experimental structure of FliC (PDB 8CXM resolves residues 3–177 and 407–498 only, so none of the mutated positions is present), the D3 domain was visualized using the AlphaFold Database model of UniProt P04949 (AF-P04949-F1), which is confidently modelled at all eight positions (pLDDT 77–90). Residues 190–283 are shown as a grey cartoon, with the side chains of the six substituted residues and the two duplication start sites drawn as sticks with Cα spheres, each individually colored and labeled with its residue index. Wild-type identities at all eight positions were verified against the full-length P04949 sequence, and the image was ray-traced in open-source PyMOL 3.1.0.

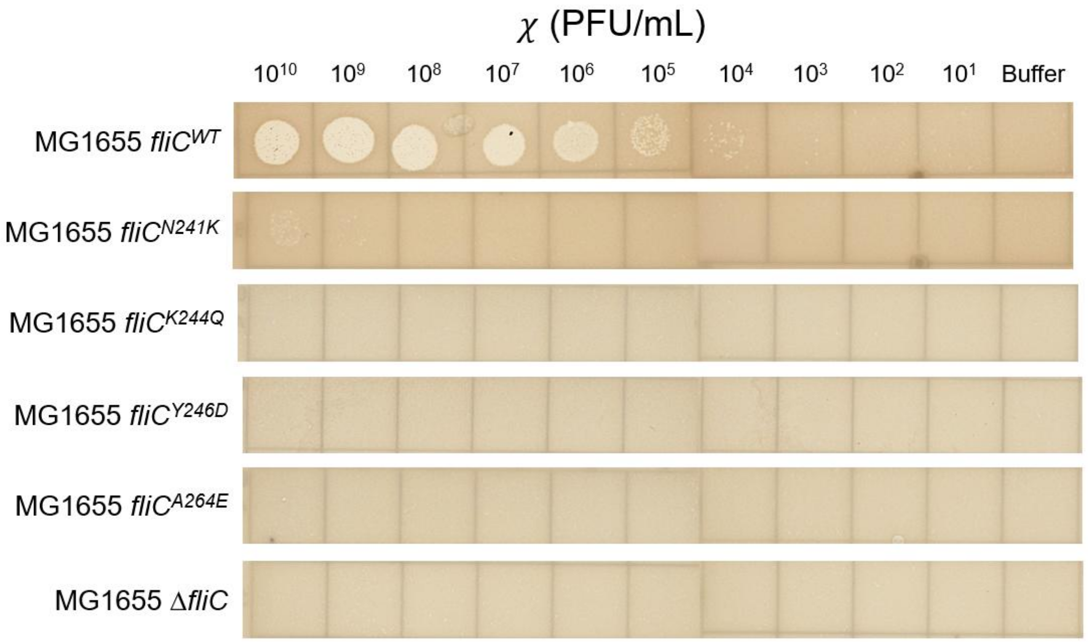


**Fig S4. Single amino acid substitutions in FliC (mutations in isolation) are sufficient to provide resistance to $\chi$.** 2.5 µL drops in a 10-fold dilution series of $\chi$ starting from $10^{10}$ PFU/mL through $10^1$ PFU/mL were deposited onto lawns of the strain indicated on the left.

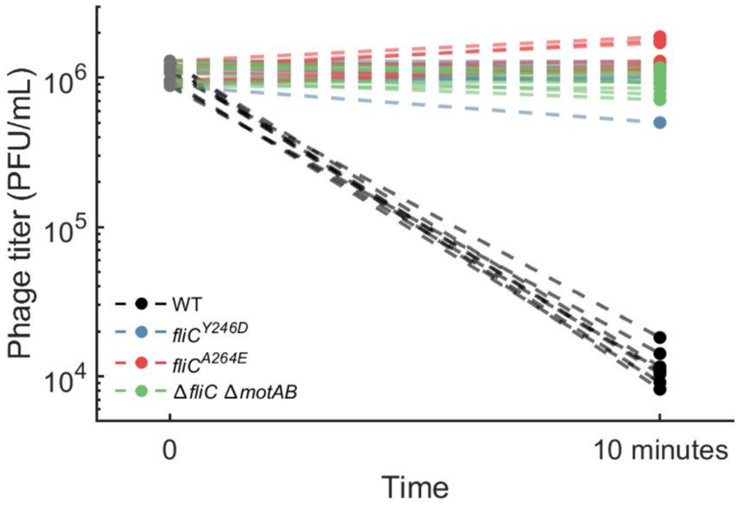


**Fig S5. $\chi$ does not adsorb to the resistant point mutants carrying amino acid substitutions in FliC.** A strain deleted for *fliC* and *motA*–*motB* that encode the flagellar stator proteins, was used as a control for lack of $\chi$ adsorption. Adsorption assays were performed using previously published protocols with some modifications [1,2].

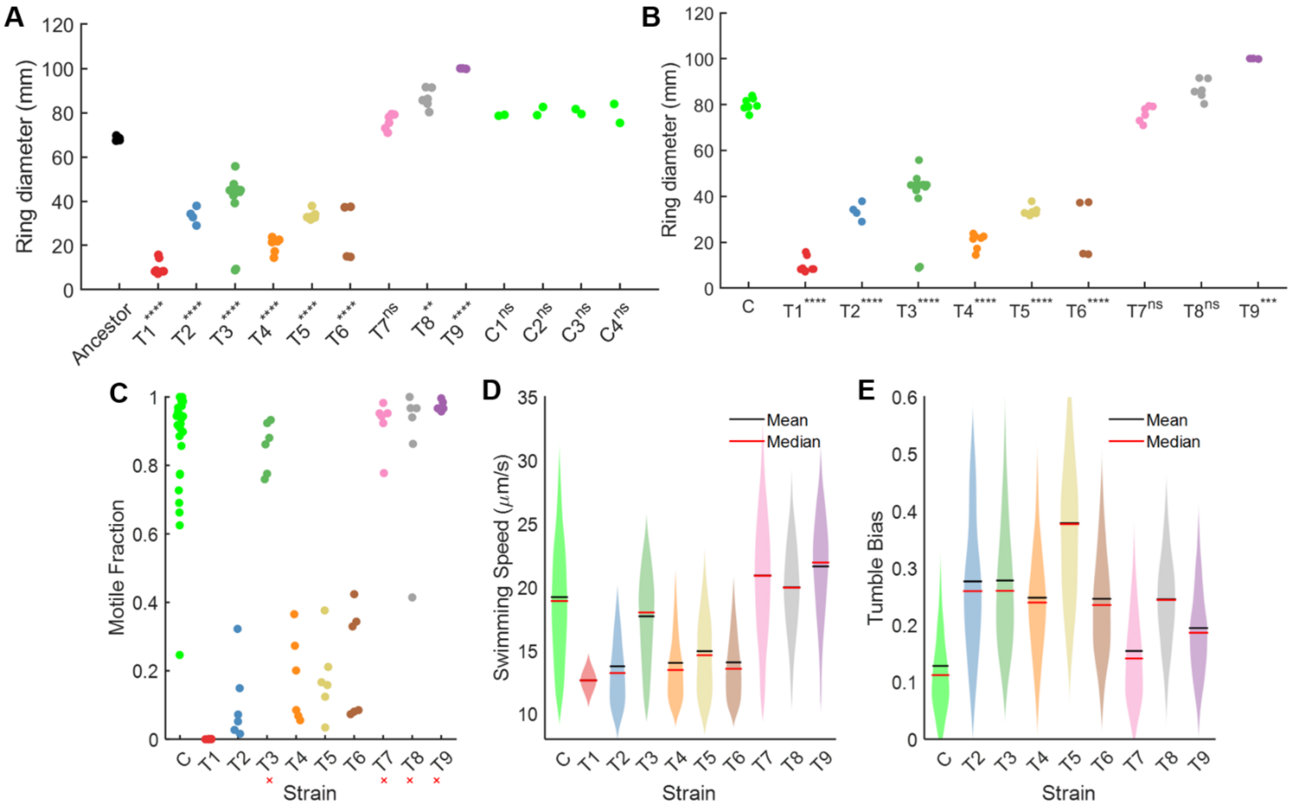


**Fig S6. Statistical tests performed for various datasets comparing Treatment strains to the Control strains.** ANOVA yielded $P < 10^{-4}$ for each test; Dunnett's post-hoc multiple hypothesis correction was applied to calculate P-values. **(A)** Swim ring diameters for strains isolated from individual lineages were compared with those of the ancestor. P-values are indicated within the X-axis titles: ** P < 0.01, **** P < 0.0001, $^{ns}$no significant difference from the Ancestor group. **(B)** Swim ring diameters for strains isolated from individual Treatment lineages were compared with those of the pooled Control (C1-C4) lineages. P-values are indicated within the X-axis titles: ** P < 0.01, **** P < 0.0001, $^{ns}$no significant difference from the Ancestor group. Motile fractions **(C)**, single-cell swimming speed distributions **(D)**, and single-cell tumble bias distributions **(E)** for each Treatment lineage were compared with those data for the pooled Control lineages. For **(C-E)**, Strains marked with a red cross (×) showed no significant difference from the pooled Controls, while all the other observations differed significantly from the pooled Controls: $P < 10^{-4}$ for all observations (not indicated). Exact P-value for each comparison is reported in **Table S4**.

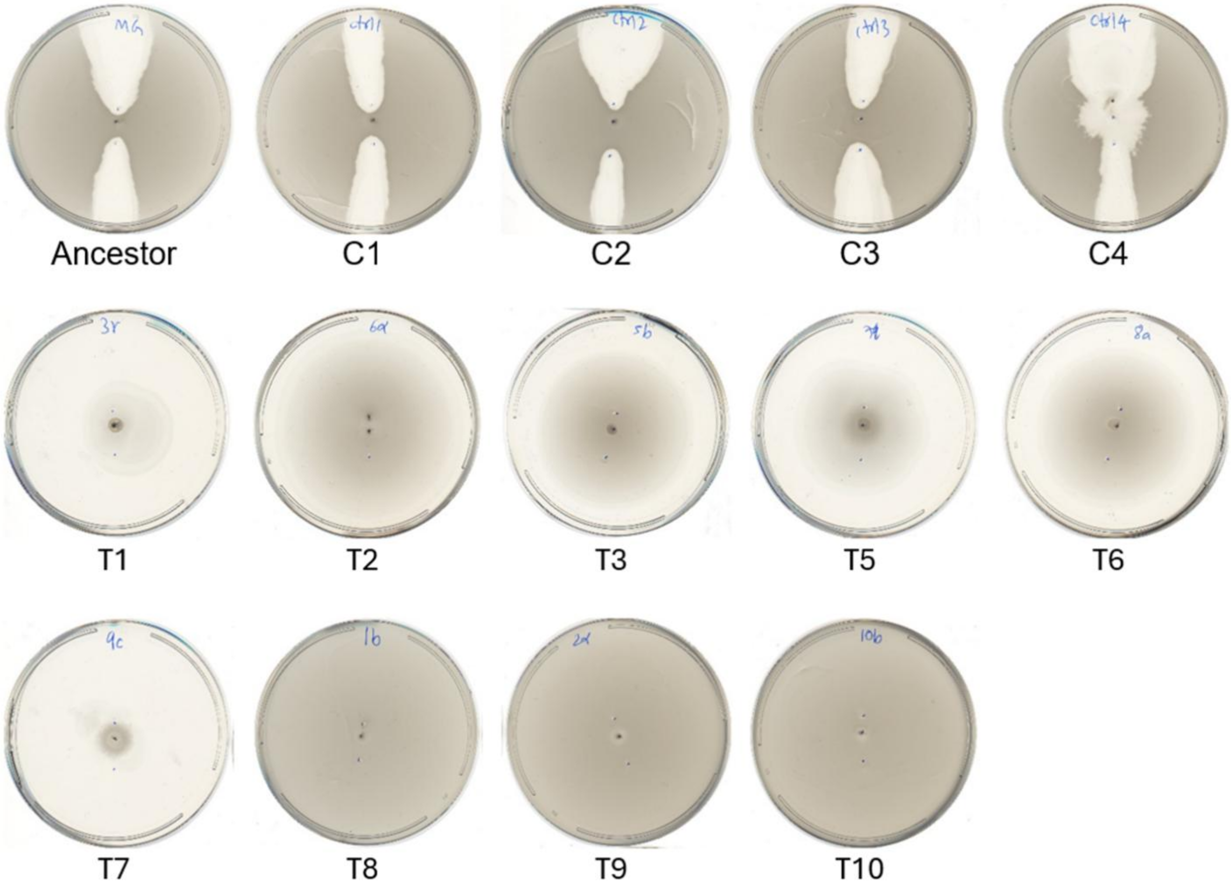


**Fig S7. Confirmation that all Treatment lineage isolates from passage 10 were motile and resistant to $\chi$, while Control lineage isolates were all motile and susceptible to $\chi$.** Highly concentrated drops of $\chi$ were deposited at the locations marked as blue dots – slightly away from the bacterial inoculum spot.

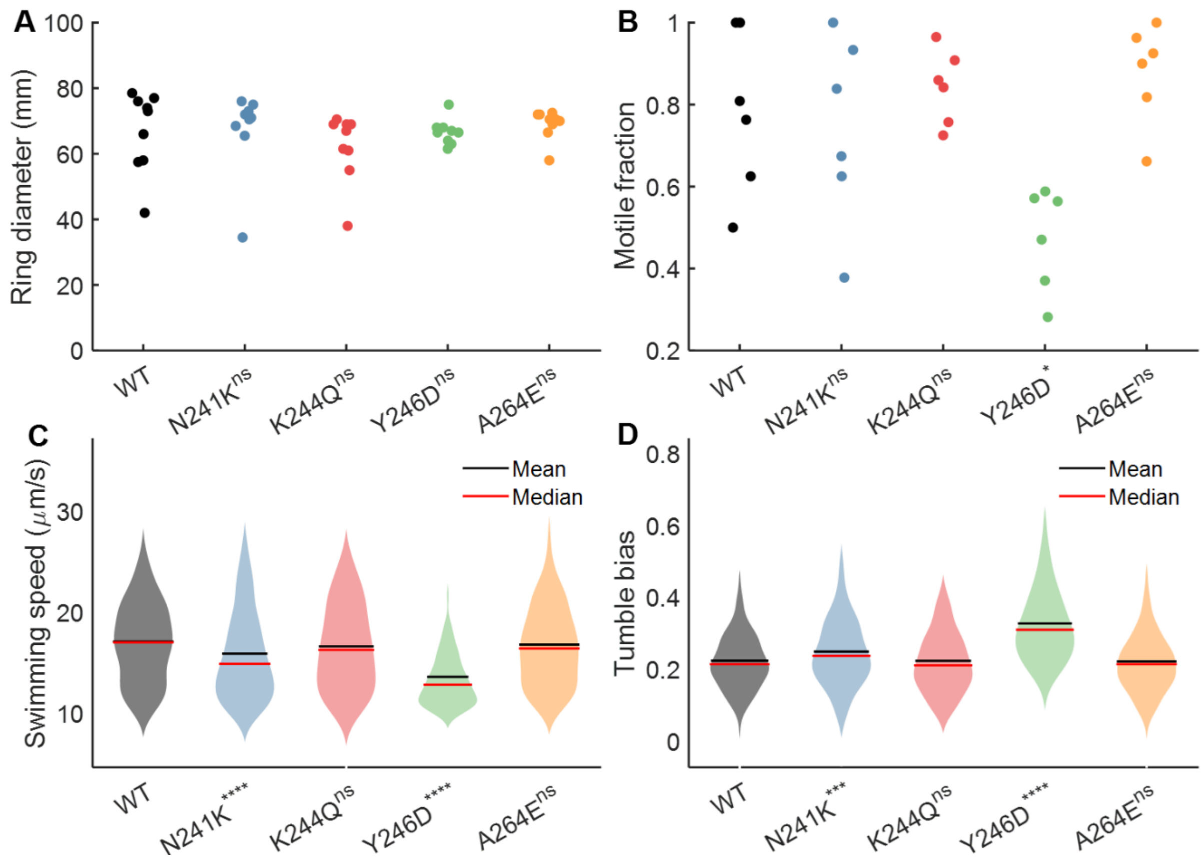


**Fig S8. Single-cell and population motility of the AA-substitution FliC mutants. (A)** Swim ring diameters of each mutant was comparable to that of WT (P = 0.65). **(B)** Motile fraction of each mutant except Y246D was comparable to that in WT (P = 0.0019). **(C, D)** Swimming speed and tumble bias of K244Q and A264E were comparable to WT, while N241K and Y246D cells had significantly reduced swimming speeds ($P < 10^{-6}$) and elevated tumble bias ($P < 10^{-6}$). P-values are reported from one-way ANOVA tests. Post-hoc Dunnett's multiple comparisons tests were performed to compare each representative lineage strain with the ancestor (* P < 0.05, *** P < 0.001, **** P < 0.0001, $^{ns}$ no significant difference from the Ancestor group).

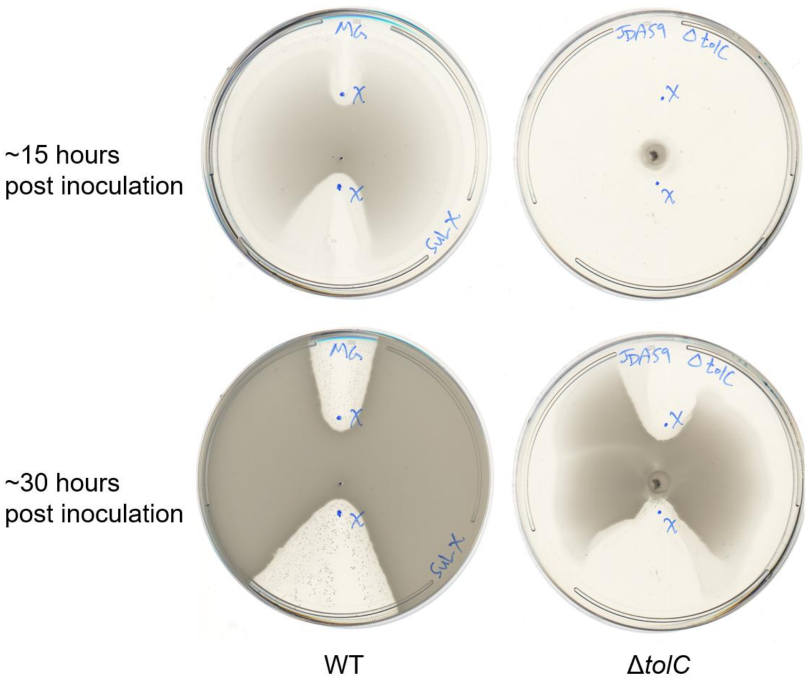


**Fig S9. Δ*tolC* cells expand slower than WT on swim plates but are susceptible to phage $\chi$.**

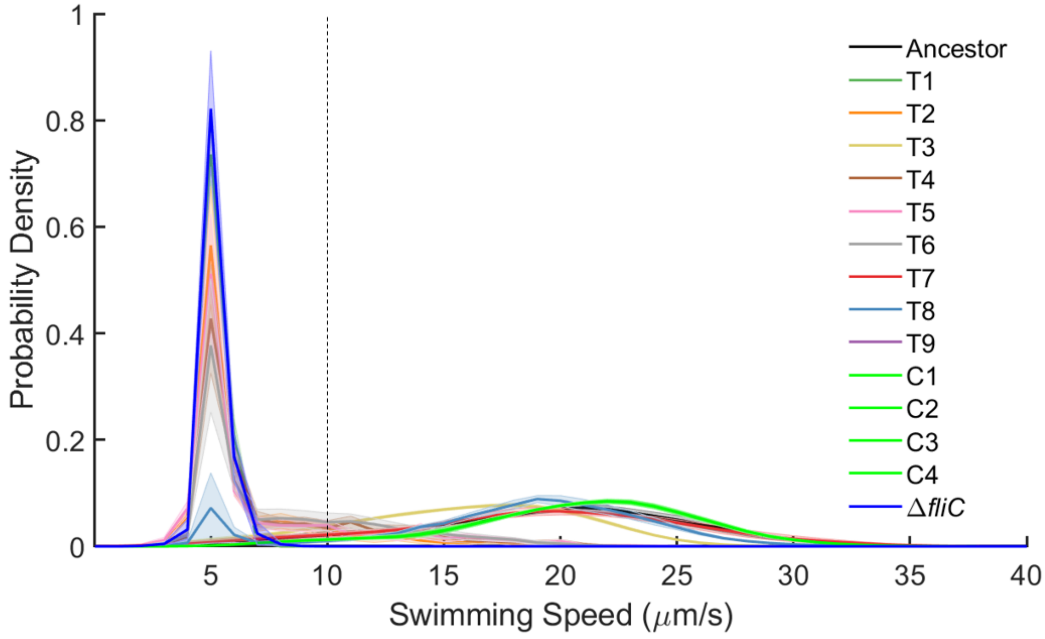


**Fig S10. Swimming speed distributions including both motile and non-motile cells.** The threshold for motile cells was determined using the MG1655 Δ*fliC::Kan* (flagellar filament knockout) strain. Based on the red distribution, we classified cells with swimming speed < 10 μm/s (left of the dashed black line) as non-motile.

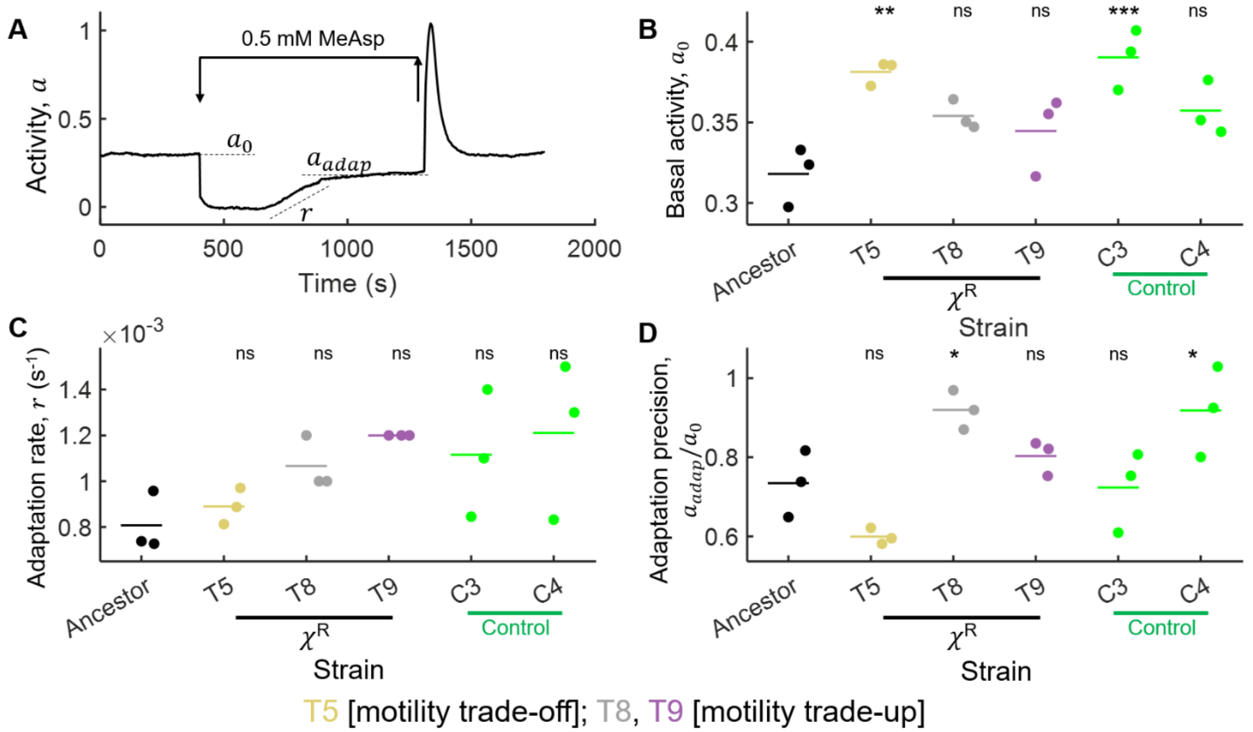


**Fig S11. FRET experiments including Controls. (A**, identical to **Fig 6**A) An example trace of the kinase activity (ratiometric FRET between CheZ-YFP and CheY-RFP) is shown. The arrows indicate the time when a saturating concentration of chemoattractant methyl-aspartate (0.5 mM) was flowed into and out of the sample chamber. Basal activity $a_0$, adapted value of activity $a_{adap}$, and adaptation rate $r$ are indicated with dashed lines ($r$ is calculated as the slope of the line, whereas $a_0$ and $a_{adap}$ are Y-values). Each trace is an output from observations of hundreds of cells within the microscopic region of interest. Basal activity $a_0$ **(B)**, adaptation rate $r$ **(C)**, and adaptation precision calculated as the ratio of the adapted and basal values of kinase activity, $a_{adap}/a_0$ **(D)** for three independent replicated experiments are indicated as dots for each strain. Mean values are indicated as dashes. ANOVA yielded P = 0.0026, 0.13, and 0.0018, respectively for **B**, **C**, and **D**. Post-hoc Dunnett's multiple comparisons tests were performed to compare each representative lineage strain with the ancestor (* P < 0.05, ** P < 0.01, ** P < 0.001, $^{ns}$ no significant difference from the Ancestor group). Exact P-value for each comparison is reported in **Table S4**.

### Supplementary Tables

**Table S1. Genes with** polymorphic sites which are over **10%** different than the ancestor sequence, **at passage 5 in Treatment lineages.** Genes known to be related to chemotaxis and flagellar motility are indicated in dark red color. Additional details are provided in S1 Data.

| Gene | Gene product |
|---|---|
| *cheB* | protein-glutamate methylesterase/protein glutamine deamidase (chemotaxis receptor demethylation enzyme) |
| *cheZ* | chemotaxis protein CheZ |
| *cysW* | sulfate/thiosulfate ABC transporter inner membrane subunit CysW |
| *dcuC* | anaerobic C4-dicarboxylate transporter DcuC |
| *dmsA* | dimethyl sulfoxide reductase subunit A |
| *drpB/rseX* | cell division protein DrpB/small regulatory RNA RseX |
| *elfC* | putative fimbrial usher protein ElfC |
| *eutD* | phosphate acetyltransferase EutD |
| *flgH* | flagellar L-ring protein |
| *fliC* | flagellar filament structural protein |
| *fliL* | flagellar protein FliL |
| *fliO* | flagellar biosynthesis protein FliO |
| *fliR* | flagellar biosynthesis protein FliR |
| *flu* | self recognizing antigen 43 (Ag43) autotransporter |
| *gyrB* | DNA gyrase subunit B |
| *hemW* | heme chaperone HemW |
| *insH8/yejO* | IS5 family transposase and trans-activator/adhesin-like autotransporter YejO |
| *kefC* | K(+) : H(+) antiporter KefC |
| *lysO/aqpZ* | L-lysine/thialysine exporter/water channel AqpZ |
| *mutL* | DNA mismatch repair protein MutL |
| *narZ* | nitrate reductase Z subunit alpha |
| *opgH* | osmoregulated periplasmic glucans biosynthesis protein H |
| *pflA* | pyruvate formate-lyase activating enzyme |
| *rhsD* | protein RhsD |
| *rnr* | RNase R |
| *rrlA* | 23S ribosomal RNA |
| *rrlG* | 23S ribosomal RNA |
| *rrsH* | 16S ribosomal RNA |
| *serS/dmsA* | serine--tRNA ligase/dimethyl sulfoxide reductase subunit A |
| *trpD* | anthranilate synthase subunit TrpD |
| *waaP* | lipopolysaccharide core heptose (I) kinase |
| *waaQ* | lipopolysaccharide core heptosyltransferase 3 |

| | |
|---|---|
| *yaiL* | DUF2058 domain-containing protein YaiL |
| *yajL* | protein/nucleic acid deglycase 3 |
| *ybhG* | HlyD_D23 family protein YbhG |
| *ycfH* | putative metal-dependent hydrolase YcfH |
| *ygaP/stpA* | thiosulfate sulfurtransferase YgaP/DNA-binding transcriptional repressor StpA with RNA chaperone activity |
| *yhhX* | putative oxidoreductase YhhX |
| *yjeT* | DUF2065 domain-containing protein YjeT |
| *ynbD* | phosphatase domain-containing protein YnbD |
| *yobD* | DUF986 domain-containing inner membrane protein YobD |
| *yohP/dusC* | putative membrane protein YohP/tRNA-dihydrouridine(16) synthase |

**Table S2. Genes with polymorphic sites which are over 10% different than the ancestor sequence, at passage 10 in Treatment lineages.** Genes known to be related to chemotaxis and flagellar motility are indicated in dark red color. Additional details are provided in S2 Data.

| **Gene** | **Gene product** |
|---|---|
| *bcp/hyfA* | thiol peroxidase/hydrogenase 4 component A |
| *cheB* | protein-glutamate methylesterase/protein glutamine deamidase (chemotaxis receptor demethylation enzyme) |
| *cheZ* | chemotaxis protein CheZ |
| *clpX* | ATP-dependent Clp protease ATP-binding subunit ClpX |
| *dnaG* | DNA primase |
| *fdnG* | formate dehydrogenase N subunit alpha |
| *fdnH* | formate dehydrogenase N subunit beta |
| *fepE* | polysaccharide co-polymerase family protein FepE |
| *flgG* | flagellar basal-body rod protein FlgG |
| *flgM/flgA* | anti-sigma factor for FliA (sigma(28))/flagellar basal body P-ring formation protein FlgA |
| *fliC* | flagellar filament structural protein |
| *fliD* | flagellar filament capping protein |
| *fliF* | flagellar basal-body MS-ring and collar protein |
| *fliG* | flagellar motor switch protein FliG |
| *fliL* | flagellar protein FliL |
| *fliM* | flagellar motor switch protein FliM |
| *fliT* | flagellar biosynthesis protein FliT |
| *ftsK* | cell division DNA translocase FtsK |
| *fumC* | fumarase C |
| *gcvT/yqfI* | aminomethyltransferase/protein YqfI |
| *glcB* | malate synthase G |
| *glgB* | 1,4-alpha-glucan branching enzyme |

| | |
|---|---|
| *gltP/yjcO* | glutamate/aspartate : H(+) symporter GltP/Sel1 repeat-containing protein YjcO |
| *hns* | DNA-binding transcriptional dual regulator H-NS (also involved in regulation of flagellar motility) |
| *hslO* | molecular chaperone Hsp33 |
| *hyfG* | hydrogenase 4 catalytic subunit HyfG |
| *kbp* | K(+) binding protein |
| *kdpD* | sensor histidine kinase KdpD |
| *lacY* | lactose permease |
| *letB* | lipophilic envelope spanning tunnel |
| *metL* | fused aspartate kinase/homoserine dehydrogenase 2 |
| *metR* | DNA-binding transcriptional dual regulator MetR |
| *mnmC* | fused tRNA 5-aminomethyl-2-thiouridylate methyltransferase and tRNA 5-aminomethyl-2-thiouridylate synthase |
| *moaA* | GTP 3',8'-cyclase |
| *motA* | motility protein A |
| *murA* | UDP-N-acetylglucosamine 1-carboxyvinyltransferase |
| *nanK* | N-acetylmannosamine kinase |
| *nuoH* | NADH:quinone oxidoreductase subunit H |
| *opgH* | osmoregulated periplasmic glucans biosynthesis protein H |
| *paaC* | phenylacetyl-CoA 1,2-epoxidase, structural subunit |
| *ppiC* | peptidyl-prolyl cis-trans isomerase C |
| *ptsP* | phosphoenolpyruvate-protein phosphotransferase PtsP |
| *radA* | DNA recombination protein |
| *rcsD* | RcsD phosphotransferase |
| *ribB/ubiK* | 3,4-dihydroxy-2-butanone-4-phosphate synthase/ubiquinone biosynthesis accessory factor UbiK |
| *rrfC* | 5S ribosomal RNA |
| *rrlA* | 23S ribosomal RNA |
| *rrlG* | 23S ribosomal RNA |
| *rrsH* | 16S ribosomal RNA |
| *tdcE* | 2-ketobutyrate formate-lyase/pyruvate formate-lyase 4 |
| *ubiI* | 2-octaprenylphenol 6-hydroxylase |
| *waaQ* | lipopolysaccharide core heptosyltransferase 3 |
| *yadI* | putative PTS enzyme IIA component YadI |
| *yaiW* | surface-exposed outer membrane lipoprotein |
| *ybaT* | putative transporter YbaT |
| *ybbP* | putative ABC transporter membrane subunit YbbP |
| *ybcV/ybcW* | DUF1398 domain-containing protein YbcV/uncharacterized protein YbcW |
| *ycbZ/matP* | putative ATP-dependent protease YcbZ/macrodomain Ter protein |
| *ycgR* | flagellar brake protein YcgR |
| *ydeQ* | putative fimbrial adhesin protein YdeQ |

| | |
|---|---|
| *yecM/argS* | putative metal-binding enzyme YecM/arginine--tRNA ligase |
| *yejK* | nucleoid-associated protein YejK |
| *ygeV* | putative sigma(54)-dependent transcriptional regulator YgeV |
| *yhaC/rnpB* | uncharacterized protein YhaC/RNase P catalytic RNA component |
| *yhbO* | protein/nucleic acid deglycase 2 |
| *yheS* | putative ATP-binding protein YheS |
| *yjjV* | putative metal-dependent hydrolase YjjV |
| *ynaM/uspF* | protein YnaM/universal stress protein F |
| *yncO/ydcC* | protein YncO/H repeat-associated putative transposase YdcC |
| *ytiE/yjiR* | protein YtiE/fused putative DNA-binding transcriptional regulator/putative aminotransferase YjiR |

**Table S3. Genes with polymorphic sites which are over 10% different than the ancestor sequence, after passage 10 in Control lineages.** Genes known to be related to chemotaxis and flagellar motility are indicated in dark red color. Additional details are provided in S3 Data.

| **Gene** | **Gene product** |
|---|---|
| *fliG* | flagellar motor switch protein FliG |
| *lpxT/mepS* | lipid A-core phosphotransferase/peptidoglycan endopeptidase/ peptidoglycan L,D-carboxypeptidase |
| *mepS* | peptidoglycan endopeptidase/peptidoglycan L,D-carboxypeptidase |
| *tsuA* | thiosulfate transporter |
| *xylE* | D-xylose:H(+) symporter |

**Table S4. Exact p-values for statistical tests.**

| **Figure** | **Data Group** | **Compared with group** | **p-value** |
|---|---|---|---|
| **Fig 5**A | T1 | Ancestor | $< 10^{-6}$ |
| **Fig 5**A | T2 | Ancestor | $< 10^{-6}$ |
| **Fig 5**A | T3 | Ancestor | 0.95 |
| **Fig 5**A | T4 | Ancestor | $< 10^{-6}$ |
| **Fig 5**A | T5 | Ancestor | $< 10^{-6}$ |
| **Fig 5**A | T6 | Ancestor | $< 10^{-6}$ |
| **Fig 5**A | T7 | Ancestor | 1 |
| **Fig 5**A | T8 | Ancestor | 0.96 |
| **Fig 5**A | T9 | Ancestor | 1 |

| | | | |
|---|---|---|---|
| **Fig 5**A | C1 | Ancestor | 1 |
| **Fig 5**A | C2 | Ancestor | 0.97 |
| **Fig 5**A | C3 | Ancestor | 0.08 |
| **Fig 5**A | C4 | Ancestor | 1 |
| **Fig 5**B | T1 | Ancestor | 0.64 |
| **Fig 5**B | T2 | Ancestor | $< 10^{-6}$ |
| **Fig 5**B | T3 | Ancestor | $< 10^{-6}$ |
| **Fig 5**B | T4 | Ancestor | $< 10^{-6}$ |
| **Fig 5**B | T5 | Ancestor | $< 10^{-6}$ |
| **Fig 5**B | T6 | Ancestor | $< 10^{-6}$ |
| **Fig 5**B | T7 | Ancestor | 0.69 |
| **Fig 5**B | T8 | Ancestor | $< 10^{-6}$ |
| **Fig 5**B | T9 | Ancestor | $< 10^{-6}$ |
| **Fig 5**B | C1 | Ancestor | $< 10^{-6}$ |
| **Fig 5**B | C2 | Ancestor | $< 10^{-6}$ |
| **Fig 5**B | C3 | Ancestor | $< 10^{-6}$ |
| **Fig 5**B | C4 | Ancestor | $< 10^{-6}$ |
| **Fig 5**C | T2 | Ancestor | $< 10^{-6}$ |
| **Fig 5**C | T3 | Ancestor | $< 10^{-6}$ |
| **Fig 5**C | T4 | Ancestor | $< 10^{-6}$ |
| **Fig 5**C | T5 | Ancestor | $< 10^{-6}$ |
| **Fig 5**C | T6 | Ancestor | $< 10^{-6}$ |
| **Fig 5**C | T7 | Ancestor | $< 10^{-6}$ |
| **Fig 5**C | T8 | Ancestor | $< 10^{-6}$ |
| **Fig 5**C | T9 | Ancestor | $< 10^{-6}$ |
| **Fig 5**C | C1 | Ancestor | $< 10^{-6}$ |
| **Fig 5**C | C2 | Ancestor | 0.0088 |
| **Fig 5**C | C3 | Ancestor | $< 10^{-6}$ |
| **Fig 5**C | C4 | Ancestor | $< 10^{-6}$ |
| **Fig 6**B | T5 | Ancestor | 0.0039 |
| **Fig 6**B | T8 | Ancestor | 0.068 |
| **Fig 6**B | T9 | Ancestor | 0.19 |
| **Fig 6**C | T5 | Ancestor | 0.61 |

| | | | |
|---|---|---|---|
| **Fig 6**C | T8 | Ancestor | 0.026 |
| **Fig 6**C | T9 | Ancestor | 0.0026 |
| **Fig 6**D | T5 | Ancestor | 0.040 |
| **Fig 6**D | T8 | Ancestor | 0.0081 |
| **Fig 6**D | T9 | Ancestor | 0.34 |
| **Fig S6**A | T1 | Ancestor | $< 10^{-6}$ |
| **Fig S6**A | T2 | Ancestor | $< 10^{-6}$ |
| **Fig S6**A | T3 | Ancestor | $< 10^{-6}$ |
| **Fig S6**A | T4 | Ancestor | $< 10^{-6}$ |
| **Fig S6**A | T5 | Ancestor | $< 10^{-6}$ |
| **Fig S6**A | T6 | Ancestor | $< 10^{-6}$ |
| **Fig S6**A | T7 | Ancestor | 0.60 |
| **Fig S6**A | T8 | Ancestor | 0.0044 |
| **Fig S6**A | T9 | Ancestor | $1.72 \times 10^{-6}$ |
| **Fig S6**A | C1 | Ancestor | 0.58 |
| **Fig S6**A | C2 | Ancestor | 0.38 |
| **Fig S6**A | C3 | Ancestor | 0.40 |
| **Fig S6**A | C4 | Ancestor | 0.49 |
| **Fig S6**B | T1 | C | $< 10^{-6}$ |
| **Fig S6**B | T2 | C | $< 10^{-6}$ |
| **Fig S6**B | T3 | C | $< 10^{-6}$ |
| **Fig S6**B | T4 | C | $< 10^{-6}$ |
| **Fig S6**B | T5 | C | $< 10^{-6}$ |
| **Fig S6**B | T6 | C | $< 10^{-6}$ |
| **Fig S6**B | T7 | C | 0.94 |
| **Fig S6**B | T8 | C | 0.53 |
| **Fig S6**B | T9 | C | $4.88 \times 10^{-4}$ |
| **Fig S6**C | T1 | C | $< 10^{-6}$ |
| **Fig S6**C | T2 | C | $< 10^{-6}$ |
| **Fig S6**C | T3 | C | 1 |
| **Fig S6**C | T4 | C | $< 10^{-6}$ |
| **Fig S6**C | T5 | C | $< 10^{-6}$ |
| **Fig S6**C | T6 | C | $< 10^{-6}$ |

| | | | |
|---|---|---|---|
| **Fig S6**C | T7 | C | 0.95 |
| **Fig S6**C | T8 | C | 1 |
| **Fig S6**C | T9 | C | 0.44 |
| **Fig S6**D | T1 | C | 0.73 |
| **Fig S6**D | T2 | C | $< 10^{-6}$ |
| **Fig S6**D | T3 | C | $< 10^{-6}$ |
| **Fig S6**D | T4 | C | $< 10^{-6}$ |
| **Fig S6**D | T5 | C | $< 10^{-6}$ |
| **Fig S6**D | T6 | C | $< 10^{-6}$ |
| **Fig S6**D | T7 | C | $< 10^{-6}$ |
| **Fig S6**D | T8 | C | $< 10^{-6}$ |
| **Fig S6**D | T9 | C | $< 10^{-6}$ |
| **Fig S6**E | T1 | C | $< 10^{-6}$ |
| **Fig S6**E | T2 | C | $< 10^{-6}$ |
| **Fig S6**E | T3 | C | $< 10^{-6}$ |
| **Fig S6**E | T4 | C | $< 10^{-6}$ |
| **Fig S6**E | T5 | C | $< 10^{-6}$ |
| **Fig S6**E | T6 | C | $< 10^{-6}$ |
| **Fig S6**E | T7 | C | $< 10^{-6}$ |
| **Fig S6**E | T8 | C | $< 10^{-6}$ |
| **Fig S6**E | T9 | C | $< 10^{-6}$ |
| **Fig S11**B | T5 | Ancestor | $2.6 \times 10^{-3}$ |
| **Fig S11**B | T8 | Ancestor | 0.085 |
| **Fig S11**B | T9 | Ancestor | 0.19 |
| **Fig S11**B | C3 | Ancestor | $9.01 \times 10^{-4}$ |
| **Fig S11**B | C4 | Ancestor | 0.056 |
| **Fig S11**C | T5 | Ancestor | 0.98 |
| **Fig S11**C | T8 | Ancestor | 0.39 |
| **Fig S11**C | T9 | Ancestor | 0.11 |
| **Fig S11**C | C3 | Ancestor | 0.26 |
| **Fig S11**C | C4 | Ancestor | 0.097 |
| **Fig S11**D | T5 | Ancestor | 0.18 |
| **Fig S11**D | T8 | Ancestor | 0.046 |

| | | | |
|---|---|---|---|
| **Fig S11**D | T9 | Ancestor | 0.72 |
| **Fig S11**D | C3 | Ancestor | 0.99 |
| **Fig S11**D | C4 | Ancestor | 0.48 |

### Supplementary note: IS1-mediated deletions

Read-depth analysis showed that IS1 remained at its native position in all six sequenced early-passage populations (**Fig 2**), while the DNA immediately adjacent to it was deleted. These deletions were not reversed in later passages. Instead, at passages 5 and 10 they persisted as subpopulations of varying frequency in 7 of 10 and 4 of 9 Treatment lineages, respectively, each with multiple, distinct deletion endpoints. They were absent from the ancestor and all Control lineages. This suggests that the motile, $\chi$-resistant populations arose from cells that never carried a *flhDC* deletion, rather than from reversion of the early nonmotile mutants. Plots indicating read depth in the relevant genomic region for each sequenced population are included in the data repository.
